# Genotypic Characterization and Antimicrobial Susceptibility of Human *Campylobacter jejuni* Isolates in Southern Spain

**DOI:** 10.1101/2024.04.17.589908

**Authors:** Pablo Fernández-Palacios, Fátima Galán-Sánchez, Carlos S Casimiro-Soriguer, Estefanía Jurado-Tarifa, Federico Arroyo, María Lara, J. Alberto Chaves, Joaquín Dopazo, Manuel A. Rodríguez-Iglesias

## Abstract

*Campylobacter jejuni* is the main cause of bacterial gastroenteritis and a public health problem worldwide. Little information is available on the genotypic characteristics of human *Campylobacter jejuni* in Spain. This study is based on an analysis of the resistome, virulome and phylogenetic relationship, antibiogram prediction and antimicrobial susceptibility of 114 human isolates of *C. jejuni* from a tertiary hospital in southern Spain from October 2020 to June 2023. The isolates were sequenced using Illumina technology, and bioinformatic analysis was subsequently performed. The susceptibility of *C. jejuni* isolates to ciprofloxacin, tetracycline, and erythromycin was tested. A high resistance rate was obtained for ciprofloxacin (90.6%) and tetracycline (66.7%), and a low resistance rate for erythromycin (0.85%) was detected among the *C. jejuni* isolates. CC-21 (n=23), ST-572 (n = 13) and ST-6532 (n=13) were the most prevalent clonal complexes (CCs) and sequence types (STs). Concerning the virulome, the *cadF, ciaB*, and *cdtABC* genes were detected in all the isolates. A prevalence of 20.1% was obtained for the genes *wlaN* and *cstIII*, which are related to the pathogenesis of Guillain-Barré syndrome (GBS). The prevalence of the main antimicrobial resistance markers detected were *cmeABC* (92.1%), *RE-cmeABC* (7.9%), the T86I substitution in *gyrA* (88.9%), *bla_OXA-61_* (72.6%)*, tet(O)* (65.8%) and *ant(6)-Ia* (17.1%). High antibiogram prediction rates (>97%) were obtained except for the erythromycin-resistant phenotype. This study contributes significantly to the knowledge of *Campylobacter jejuni* genomics for the prevention, treatment and control of infections caused by this pathogen, which is relevant to public health.

**Importance:** Despite being the pathogen with the greatest number of gastroenteritis cases worldwide, *Campylobacter jejuni* remains a poorly studied microorganism. The development of whole-genome sequencing (WGS) techniques has led to a better understanding of the genotypic characteristics of this pathogen. These techniques complement the data obtained from the phenotypic analysis of *C. jejuni* isolates. The zoonotic transmission of *C. jejuni* through the consumption of contaminated poultry implies approaching the study of this pathogen through the term “One Health.” This is the first study, using WGS, conducted on human isolates of *C. jejuni* in Spain to date, which allows comparison of the results obtained with similar studies conducted in other countries and with animal and environmental isolates.

## Introduction

*Campylobacter* is the main cause of bacterial diarrheal disease, with over 95 million cases reported and 20,000 deaths worldwide (1). Although infections are usually self-limiting, *Campylobacter* can also cause severe infections such as bacteremia, sepsis, arthritis, endocarditis, or Guillain–Barré syndrome (GBS) (2). *Campylobacter* is part of the WHO list of priority pathogens for the research and development of new antibiotics due to its high resistance to fluoroquinolones in human isolates (3). Spain has the highest combined resistance rate to ciprofloxacin and erythromycin (3.1%) among the human isolates of *C. jejuni* in Europe (4).

Point mutations in the *gyrA* gene are the principal mechanism of resistance in *Campylobacter* (5). In tetracyclines, antibiotic resistance is generated by the presence of the *tet(O)* gene. In macrolides, resistance is caused by mutations in the 23S ribosomal RNA gene and by the presence of the *erm(B)* gene, which encodes a methylase. Resistance to beta-lactams in *Campylobacter* is mainly due to the beta-lactamase OXA-61 encoded by the *bla_OXA-61_ gene* (6,7). Several studies have revealed the presence of genes that encode aminoglycoside-modifying enzymes, such as *the ant*(*6*)*-Ia* and *aph(3’)-III* genes, in *Campylobacter* (8, 9). In addition, antibiotic efflux pumps (CmeABC, CmeFED, and CmeG) contribute to antibiotic resistance (10). Furthermore, the presence of the resistance-enhancing variant CmeABC (named RE-CmeABC), which significantly increases resistance to ciprofloxacin and erythromycin, has been described in human isolates of *C. jejuni* (11).

The mechanisms underlying the pathogenicity and virulence of *C. jejuni* are not fully understood. However, several genes involved in motility, cell cytotoxicity, invasion, chemotaxis, and the cellular stress response have been described (12–14). Virulence factors *(wlaN, cgtB, cstII,* and *cstIII)* implicated in the pathogenesis of GBS via the modification of lipooligosaccharide (LOS) have also been studied (15, 16).

Whole-genome sequencing (WGS) has allowed significant advances in the field of molecular epidemiology using the cgMLST (multilocus core genome sequence typing) and wgMLST (whole genome multilocus sequence typing) schemes (17). Both have greater discriminatory power than the traditional MLST scheme (18).

To date, there is little information available on the WGS data obtained from *Campylobacter* clinical isolates in Europe (19–21) and Spain. The objective of this study was to analyze the phenotypic and genotypic characteristics of 114 clinical *C. jejuni* isolates from southern Spain. Advanced WGS tools enable a more in-depth understanding of the resistome, virulome, and epidemiology. In this way, we can better understand the pathogenicity and nature of *C. jejuni*, a highly important zoonotic pathogen for public health.

## Methods

### Isolates

A total of 114 *C. jejuni* isolates recovered from unformed stool from patients with diarrhea submitted to the microbiology laboratory of a tertiary hospital in southern Spain from October 2020 to June 2023 were included in the study. The patients came from the cities of Cadiz and San Fernando. The samples were cultured in blood-free BD Campylobacter selective medium (Beckton Dickinson, New Jersey, USA) and incubated at 42 °C for 48 hours under microaerobic conditions (85% nitrogen, 10% carbon dioxide, and 5% oxygen). The *C. jejuni* isolates were confirmed by mass spectrometry using MALDI-TOF (Bruker, Bremen, Germany).

### Antimicrobial susceptibility testing

Antimicrobial susceptibility testing was performed using the disk diffusion method according to the European Committee on Antimicrobial Susceptibility Testing (EUCAST) recommendations (22). The antibiotics tested included ciprofloxacin (5 μg), erythromycin (15 μg) and tetracycline (30 μg) (Oxoid, Basingstoke, UK). The isolates were incubated for 24-48 hours under microaerophilic conditions, and the zones of antibiotic inhibition were interpreted according to EUCAST breakpoints 2023 (22).

### Whole-genome sequencing and data analysis

Bacterial DNA was extracted from *C. jejuni* isolates using the EZ1 Advanced XL (Qiagen, Hilden, Germany) following the manufacturer’s instructions. The quantity of extracted bacterial DNA was measured using the Qubit 3.0 fluorometer (Fisher Scientific, Massachusetts, USA) and the Qubit dsDNA HS assay kit. Libraries were prepared using the Nextera XT DNA Library Preparation Kit (Illumina, Cambridge, UK) according to the manufacturer’s instructions and quantified using the Qubit 3.0 fluorometer. MiSeq (Illumina, Cambridge, UK) was used for the genome sequencing of the isolates. Quast (23), Kraken (24), Fastp (25), Qualimap (26), and Checkm (27) were used to evaluate the quality of the FASTQ files obtained in the sequencing process and the FASTA files of the assembled genomes. De novo assembly was performed using SPAdes (28). The data obtained from the use of these bioinformatics tools and those obtained through the analysis of *C. jejuni* sequences are deposited in the SIEGA app (Andalusia Integrated Molecular Epidemiology System) (29). The genome metrics of the isolates are available in Table S1 in the supplemental material.

### Clonal complex (CC), multilocus sequence typing (MLST), and core genome MLST (cgMLST) analyses

The MentaLiST application (30) was used to obtain allele profiles from *C. jejuni* isolates. Based on these allelic profiles, the sequence types (STs) and CCs of the isolates were determined using the MLST scheme, which is based on the variation in seven highly conserved genes (*aspA, glnA, gltA, glyA, pgM, tkt,* and *uncA*). The core genome MLST (cgMLST) analysis of the isolates was conducted using allelic profiles based on 1,343 core genome genes of *Campylobacte*r (31) sourced from the PubMLST database. The *C. jejuni*/*C. coli* cgMLST v1 version, updated as of January 5, 2024, was obtained from PubMLST. The alleles were identified using an in-house developed script. A neighbor-joining phylogenetic tree was constructed to establish the phylogenetic relationships among the isolates according to the cgMLST scheme. The phylogenetic results and annotations were visualized using iTOL v.6 (32).

### Identification of virulence factors

The Virulence Factor Database (VFDB) (33) was used to determine the virulence factors present in the *C. jejuni* isolates. From the total virulence genes obtained by VFBD, *cadF* (cell adhesion), *ciaB* (invasion) and *cdtABC* (cytotoxin production) were analyzed. The presence of genes related to GBS pathogenesis that modify the structure of lipopolysaccharides (LOS), such as *wlaN* and *cstIII*, and genes related to type IV secretion system (T4SS), such as *VirB* genes, was also investigated.

### Identification of antimicrobial resistance markers and correlation between phenotype and genotype

Antimicrobial resistance markers (*tet(O)* gene, *erm(B)* gene, aminoglycoside resistance genes, blaOXA*_-61_* gene, and *cmeABC* operon) present in the *C. jejuni* sequences were obtained using the ABRicate pipeline (34). The databases used were Resfinder (35), CARD (36), MEGARes (37) and ARG-ANNOT (38). Chromosomal mutations in the *gyrA* and 23S rRNA genes were obtained using PointFinder (39), which is included in Resfinder version 4.4.2. Correlations between the resistance phenotypes to ciprofloxacin, erythromycin, and tetracycline and the genotypes (antimicrobial resistance markers) of 114 *C. jejuni* isolates were determined. The BLASTn tool (40) was used to search for the presence of RE-CmeABC in isolates where the typical *cmeB* subunit was not detected.

### Accession number

The sequences used for the analysis of RE-CmeABC were RE-CmeABC (GenBank genome accession: KT778507.1) and the *cmeB* subunit (GenBank genome accession: NC_002163.1). Full data on antimicrobial resistance markers, virulence factors, antimicrobial susceptibility testing, and genetic diversity are available in the supplementary files.

## Results

### Phenotypic results of antimicrobial susceptibility testing

Antibiotic susceptibility tests were performed on 114 isolates of *C. jejuni* (see Table S2 in the supplemental material). All results were interpreted at 24 hours according to EUCAST breakpoints. The resistance rates for each antibiotic were 90.3% for ciprofloxacin, 66.7% for tetracycline and 0.88% for erythromycin. Twenty-eight isolates (24.6%) were resistant to at least one antibiotic, while 77 isolates (67.5%) were resistant to at least two antibiotics. The phenotype resistant to ciprofloxacin and tetracycline (65%) was the most common among the *C. jejuni* isolates. Nine isolates (7.9%) were susceptible to the three antibiotics. Only one isolate (cjeju22_21) was resistant to erythromycin. Table 1 shows the antibiotic resistance data and the number of antibiotics used for each observed phenotype.

**Table 1.** Antimicrobial susceptibility testing was performed on 114 isolates of *C. jejuni.* The antibiotics tested were ciprofloxacin (CIP), tetracycline (TET), and erythromycin (E).

| Number of resistant antimicrobial group | Phenotype (CIP, TET, E) | Number of isolates (%) |
| --- | --- | --- |
| 0 | (S,S,S) | 9 (7.9) |
| 1 | (R,S,S) | 28 (24.6) |
| 1 | (S,R,S) | 2 (1.7) |
| 2 | (R,R,S) | 74 (64.9) |
| 2 | (R,S,R) | 1 (0.9) |

### Phylogenetic study of *Campylobacter jejuni* isolates

A variety of 14 CCs, 41 STs, and 68 different cgSTs were obtained among the 114 isolates of *C. jejuni* (see Table S3 in the supplemental material). CC-21 (n=23), CC-206 (n=17), and CC-353 (n=14) were the most prevalent CCs. ST-572 (n=13), ST-6532 (n=13), and ST-50 (n=10) were the most prevalent STs. Among the wide variety of cgTs detected, the most prevalent were cg85207 (4 isolates distributed across two clusters of two cases each between 2022 and 2023), cg78144 (4 isolates distributed across two clusters of two cases each in 2022), cg35104 (4 isolates distributed across two clusters of two cases each between 2020 and 2021) and cg19242 (4 isolates distributed across two clusters of two cases each between 2020 and 2021) (Figure 1).

**Figure 1.**
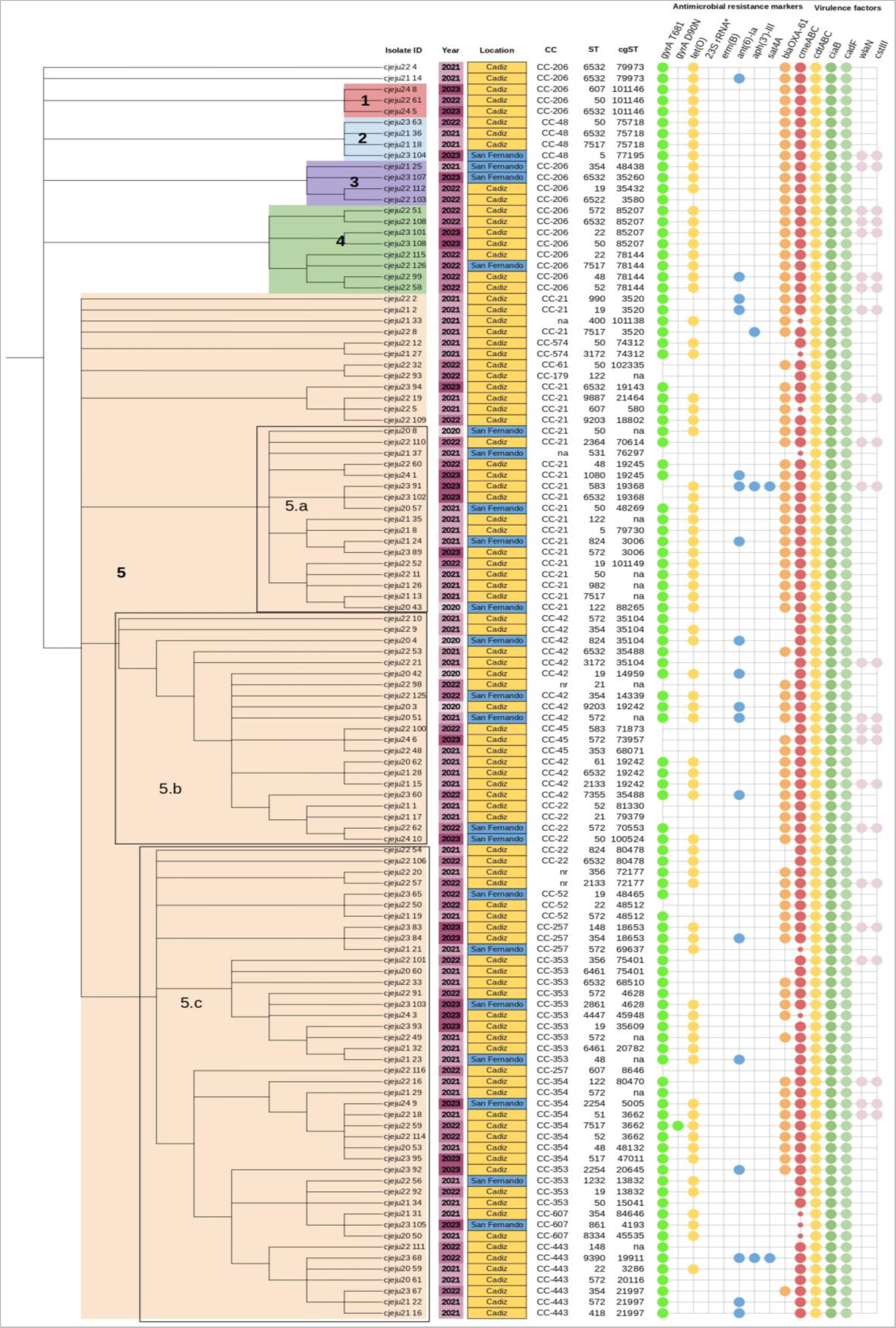
Phylogenetic analysis of 114 isolates of *C. jejuni.* The phylogenetic tree is based on the cgMLST scheme and visualized by iTOl v.6. The clades are denoted by the numbers 1, 2, 3, 4 and 5. Clade 5 is divided into subclades 5.1, 5.2, and 5.3. Clonal complexes (CCs), sequence types (STs), and core genome sequence types (cgSTs) are listed on the labels of each isolate. The presence of antimicrobial resistance at markers and virulence factors is visualized by circles in colors for each gene. Location (Cadiz and San Fernando) and year of isolation (from 2021 to 2023) are represented in colors. The isolates that could not be assigned a single cgST were labeled ’na’ (not assigned). The presence of *RE-cmeABC* operon is indicated by a small red circle. (*) 23S rRNA mutations.

Five isolates (4.4%) did not belong to the described CCs but were included in 4 STs: ST-2133 (n=2), ST-2861 (n = 1), ST-1080 (n=1) and ST-531 (n=1). For 12 isolates, an exact cgMLST profile could not be assigned; more than one cgMLST profile was detected according to *C. jejuni/C. coli* cgMLST scheme (see Table S3 in the supplemental material). The CCs, STs, and cgSTs of each isolate are shown in Figure 1.

Phylogenetic analysis revealed that the *C. jejuni* isolates clustered into 5 different clades. Two isolates (cjeju22_4 and cjeju21_14) did not cluster into any clade (Figure 1). Clade 1 is composed of 3 isolates recovered in Cadiz in the same year and shares the same CC (CC-206) and cgST (cg101146), although they are distributed in three STs (Figure 1). Clade 2 and clade 3 were each composed of 4 isolates. Isolates belonging to these clades were recovered in Cadiz and San Fernando between 2021 and 2023 and share the same CC among them, CC-206 for the isolates of clade 2 and CC-48 for clade 3. However, isolates belonging to clade 2 and clade 3 were grouped into different STs and cgSTs (Figure 1). Clade 4 was composed of 8 isolates, 7 of which were recovered in Cadiz and one in San Fernando between 2022 and 2023. All isolates in this clade belong to CC-206, and they are clustered into 7 STs and 2 cgSTs (cg85207 and cg78144) (Figure 1). Clade 5 was composed of 93 isolates, which were further divided into 3 subclades (5.1, 5.2, and 5.3). Of the 93 isolates belonging to clade 5, 12 did not belong to any subclade (Figure 1). Isolates belonging to clade 5 exhibit great variability according to the year of isolation, geographical distribution and epidemiology. None of the five clades showed a clear distribution of any specific marker of antimicrobial resistance.

### Virulence factors

A wide variety of virulence factors were detected in the *C. jejuni* isolates (Figure 1). The most prevalent virulence factors were associated with cell adhesion (*cadF*), invasion (*ciaB*), and cytotoxin production (*cdtABC*) and were detected in all the *C. jejuni isolates*. The *wlaN* and *cstIII* genes, which are implicated in the pathogenesis of GBS, were detected in 23 isolates (20.15%). These 23 isolates were distributed in 7 CCs (Figure 1). The *wlaN* and *cstIII* genes were detected mainly in CC-21 (n = 13) and CC-206 (n=4). Three out of four isolates belonged to the cg85207 group, which carries the *wlaN* and *cstIII* genes (Figure 1). Genes related to type 4 secretion system (T4SS) were not detected in any isolate of *C. jejuni*.

### Antimicrobial resistance markers

*GyrA* mutations were detected in 101 isolates (88.6%). The T86I substitution was detected in these 101 isolates. In addition, the D90N substitution was detected in isolate cjeju22_59. The *tet(O)* gene was identified in 75 isolates (65.8%). Regarding aminoglycoside resistance genes, the *ant*(*6*)*-Ia* gene was detected in 19 isolates (16.7%). In the isolates cjeju23_68 and cjeju23_91, the *ant*(*6*)*-Ia* gene, *aph(3’)-III* gene, and *sat-4* gene were detected (Figure 1). The *aph(3’)-III* gene encodes a phosphotransferase that confers resistance to kanamycin, amikacin, and other aminoglycosides. The *sat-4* gene encodes resistance to streptomycin. Furthermore, *the aph(3’)-III* gene was detected in the isolate cjeju22_8 (Figure 1).

A high distribution of the *cmeABC* operon was detected between the 105 *C. jejuni* isolates (92.1%). In 9 isolates (7.9%), the *cmeA* and *cmeC* subunits but not the *cmeB* subunit were detected (Figure 1). In these 9 isolates, the presence of RE-CmeABC was detected (95% identity) using the BLASTn tool, leading to the hypothesis that these isolates carry RE-CmeABC. The *bla_OXA-61_* gene was detected in 83 isolates (72.8%). Point mutations in the 23S rRNA and *erm(B)* genes were not detected in any of the isolates. All resistance markers detected in the *C. jejuni* isolates are shown in Figure 1.

### Correlation between resistance phenotype and antimicrobial resistance markers

Correlations were determined between antimicrobial susceptibility test (phenotype) and WGS (genotype) data (see Table S4 in the supplemental material). The substitution of T68I in the *gyrA* gene was detected in 102 isolates resistant to ciprofloxacin but was not detected in any of the isolates susceptible to ciprofloxacin, with correlations of 98.1% and 100%, respectively. Seventy-three out of the seventy-six isolates resistant to tetracycline carried the *tet(O)* gene, indicating a correlation between the phenotype and genotype of 97.4%. Similarly, none of the isolates susceptible to tetracycline carried the *tet(O)* gene, showing a 100% correlation. Resistance markers for erythromycin resistance were not detected among the susceptible isolates, indicating a 100% correlation between phenotype and genotype. One isolate was resistant to erythromycin, but neither the *erm(B)* gene nor any mutations in 23S rRNA were detected. The correlations between the phenotypes and genotypes of *C. jejuni* isolates treated with ciprofloxacin, tetracycline and erythromycin are presented in Table 2.

**Table 2.** Correlation between the phenotype and genotype of the 114 isolates of *C. jejuni*. Antimicrobial resistance markers detected for each clinical category for ciprofloxacin, tetracycline and erythromycin are shown. The isolates were classified by clinical category according to the EUCAST breakpoints: resistant (R) and susceptible standard dosing (S) regimens.

| Antibiotic | Number of isolates R o S; | Number of isolates carrying resistance markers | Rate of correlation between phenotype/genotype (%) |
| --- | --- | --- | --- |
| Ciprofloxacin | R: 103 | 101 (*gyrA* T86I)<br>1 (*gyrA* T86I+ *gyrA* D90N) | 98.1 (101/103) |
|  | S: 11 | 0 | 100 (11/11) |
| Erythromycin | R: 1 | 0 | 0 |
|  | S: 113 | 0 | 100 (113/113) |
| Tetracycline | R: 76 | 74 (*tet(O)*) | 97.4 (74/76) |
|  | S: 38 | 1 (*tet(O)*) | 97.4 (37/38) |

## Discussion

This analysis, performed on clinical isolates of *C. jejuni,* included the study of the resistome and virulome and phylogenetic analysis using whole-genome sequencing, as well as phenotypic antimicrobial susceptibility data. To the best of our knowledge, this is the first study of these characteristics in human isolates of *Campylobacter jejuni* in Spain.

Ciprofloxacin, doxycycline, and azithromycin are the antibiotics most commonly used for the treatment of Campylobacter-related gastroenteritis (2). Due to its low resistance rate, azithromycin is considered the antibiotic of choice. The resistance rates for ciprofloxacin and tetracycline determined in this study (90.3% and 66.7%, respectively) are higher than the European averages of 69.1% and 46.6%, respectively (4), but are similar to those reported for Spanish isolates of *C. jejuni* in 2022 (41).

Phylogenetic analysis revealed that CC-21 is the most common clonal complex in our area (20.2%). This finding is consistent with genomic studies conducted in South America (42–44) and Europe (19–21). In a study that included 108 Peruvian isolates of *C. jejuni,* CC-362 was the most prevalent clonal complex, followed by CC-21 (44). In addition to CC-21, Spanish isolates were grouped mainly into CC-206 (11.4%) and CC-353 (11.4%), with significantly greater percentages than those of PubMLST (5.3% and 5.5%, respectively) (45).

Among the varieties of STs obtained, 10 (8.8%) were ST-50. ST-50 is widely distributed around the world and is detected in greater or lesser proportions depending on the geographical area (19–21, 42). In this study, the most common STs were ST-572 (11.4%) and ST-6532 (11.4%). According to the data of clinical isolates uploaded to PubMLST (45), 2 of the 103 Spanish isolates were ST-572, and none were ST-6532. ST-572 has been detected almost entirely in Europe in animal and human isolates. According to PubMLST (45) ST-6532 is less widely distributed than ST-572 and is detected in 15 isolates, most of which are in Great Britain. The results of this analysis of isolates from southern Spain are partially in agreement with those obtained in other European countries with respect to the distribution of ST-572 and ST-50 (45). However, in this geographic area, a greater distribution of ST-6352 was detected in the Spanish isolates. ST-6532 was detected in 3 isolates from a genomic study of 40 *C. jejuni* isolates from ruminants in Vizcaya (Northern Spain) (46). Analysis of the cgMLST profiles of the isolates revealed a heterogeneous distribution without a clear predominance of any specific cgST. Cg35104, cg19242, cg85207 and cg78144 were the most prevalent, although not in a large proportion of the total isolates.

The isolates in our study were grouped into 5 different clades with high genetic diversity. Regarding the study of phylogenetic clades, previous studies in Peru and Poland have shown that associations between clusters and the presence of antimicrobial resistance markers and virulence factors are common (44, 47). In our study, we did not observe an epidemiological relationship in this regard. Furthermore, there was no observable distribution based on geographic area, Cadiz or San Fernando (Figure 1), as these populations are very close, and there was a significant interaction between the populations of both cities.

The mechanisms of antibiotic resistance in *Campylobacter* have been widely studied. However, the pathogenic mechanisms and the mode of interaction of *Campylobacter* with the host are not fully defined (12, 48). A prevalence of 21% was detected for the *wlaN* and *cstIII* genes, with a possible association with ST-50 because these two genes were detected in 80% of the isolates belonging to ST-50 (Figure 1). The prevalence of the *wlaN* gene is similar to that obtained in a study conducted in northern Spain, where the *wlaN* gene was detected in 20% of human isolates. However, there is no clear predominance of any ST that could indicate a relationship between ST and the distribution of this gene (15).

The C257T mutation, which causes the substitution of T68I in the *gyrA* gene, is the main mechanism of resistance to ciprofloxacin reported in several studies (19, 20, 42, 43). In our study, the T68I substitution was detected in almost 90% of the isolates (Figure 1). Other mutations in the *gyrA* gene that also confer resistance to ciprofloxacin have been described, although they are detected in smaller proportions, such as D90N and T86K (49). In the isolate cjeju22_59, the D90N substitution was detected along with the T86I substitution. The percentage of detection of the *tet(O)* gene, *bla_OXA-61_*, and *cmeABC* operon aligns with the findings of previous studies conducted in other countries (20, 42, 43). The main gene encoding aminoglycoside resistance genes was *ant*(*6*)*-Ia* (16.7%), which confers resistance to streptomycin. This gene was detected in the same proportion in studies performed in North America and South America in *C. jejuni* clinical isolates (50). The *aph(3’)-III* gene was detected in three isolates (cjeju23_92, cjeju22_49 and cjeju24_3). In isolates cjeju22_49 and cjeju24_3, *t*he *aph(3’)-III* gene was detected along with the *ant*(*6*)*-Ia* gene and the *sat-4* gene. These three genes can be included in these two isolates, forming the *(ant*(*6*)*-sat4-aph(3′)-III*) cluster. This cluster has been detected in the chromosomal and plasmidic genomes of gram-positive bacteria such as *Streptococcus* (51). No antimicrobial resistance markers for erythromycin or type IV secretion (T4SS) were detected, which is consistent with the low prevalence of these genes in clinical isolates of *C. jejuni* (20, 42, 52).

Another interesting result of our study is the detection of the RE-CmeABC variant in human *C. jejuni* isolates in Spain. The fact that antibiotic resistance databases do not include the sequence of this efflux pump and the few published studies to date about the characteristics of RE-CmeABC have resulted in a lack of information about this resistance mechanism in *C. jejuni.* The presence of this RE-CmeABC variant can be suspected when the *cmeB* subunit from the CARD (53) is not detected, and atypical *cmeB* can be detected by comparing the *cmeB* subunit of *C. jejuni* NCTC 1168 with that of an isolate (11). A recent study conducted in Peru on 97 clinical isolates of *C. jejuni* indicated a prevalence of 36.1% for RE-CmeABC (53). In our study, this variant was detected in 8% of the isolates, 5 of which belonged to subclade 5.3. Three of these isolates exhibited a phylogenetic relationship. These three isolates belong to CC-607, but no epidemiological relationship is observed according to the geographic area and year of isolation (Figure 1).

WGS-based antibiotic resistance tools have yielded excellent results in predicting antibiogram outcomes. A correlation of more than 98% was obtained between antimicrobial resistance markers and antimicrobial susceptibility tests for ciprofloxacin, tetracycline and erythromycin among more than 500 isolates of *C. jejuni* in Denmark (54). Other analyses performed in Israel revealed 96-100% correlation among 263 isolates of *C. jejuni* (55). These results are similar to those obtained in our analysis of 114 isolates, as more than 97% correlation was obtained (except for the erythromycin-resistant phenotype). According to these previous studies, tetracycline is the antibiotic with the lowest correlation percentage (54,55). Other resistance mechanisms, such as the overexpression of the CmeABC efflux pump, may explain why any antimicrobial resistance genes related to these antibiotics were detected in the isolates resistant to ciprofloxacin, tetracycline, and erythromycin (56–58).

A limitation of this analysis is that the collection of *C. jejuni* isolates was not carried out through a continuous surveillance programme but rather from a specific period. This may not accurately represent the characteristics of *C. jejuni* in our region. Furthermore, 4.4% of the isolates could not be typed by CC, and 10.5% of the isolates could not be typed by cgMLST, providing a loss of relevant information for understanding the local epidemiology of isolates. Another limitation of this study is that the isolates were recovered from a small geographical area in southern Spain.

In conclusion, our study provides new data about the genotypic and phenotypic characteristics of human *C. jejuni* isolates from Spain. The results obtained in this work have been deposited in the database of the Integrated Genomic Surveillance System of Andalusia (SIEGA). The SIEGA project was created with the objective of obtaining, through WGS, a genomic database of the most important pathogens for public health in Andalusia. The findings obtained in this work should be correlated with data on animal and environmental isolates, providing data about the relationship between animal, human, and environmental health under the “One Health” approach.

## Acknowledgments

This work was supported by the Integrated Genomic Surveillance System of Andalusia (SIEGA), Fundación Progreso y Salud, Consejería de Salud y Consumo, Junta de Andalucía, Spain. CSCS was funded by a Juan de la Cierva grant (FJC2021-046546-I) from the Ministerio de Ciencia e Innovación.

## Data availability

The sequences of all the isolates sequenced in this study are available at GenBank (project number PRJNA1088320, https://dataview.ncbi.nlm.nih.gov/?archive=bioproject). The metadata of the isolates and MLST profiles are available at PubMLST (https://pubmlst.org/bigsdb?db=pubmlst_campylobacter_isolates).

## Supplemental material

Table 1S

Table 2S

Table 3S

Table 4S

PFP, FGS, and MRI contributed to the study conception and design. PFP and FA were responsible for conducting the antibiotic susceptibility test. EJT, CSCS, MA, and JD performed the sequencing and bioinformatic analysis with the support of JACS. The first draft of the manuscript was written by PFP, FGS, and MRI, and all the authors commented on previous versions of the manuscript. All the authors have read and approved the final manuscript.

## Compliance with ethical standards

**Conflict of interest**

**Ethics approval** Not applicable.

**Consent to participate** Not applicable.

**Consent for publication** Not applicable

**Financial Interest** The authors declare that they have no financial interests.

---

**Table S1.** Genome metrics and accession numbers of 114 *Campylobacter jejuni* isolates.

| **SRA accession** | **Genome accession** | **Biosample_accession** | **Isolate ID** | **Genome coverage** | **N50** | **Number of reads** | **Number of contigs** |
| --- | --- | --- | --- | --- | --- | --- | --- |
| SRR28623923 | JBBFGA000000000 | SAMN40466906 | cjeju20_3 | 99.98x | 139291 | 5734618 | 40 |
| SRR28623922 | JBBFFZ000000000 | SAMN40466907 | cjeju20_4 | 99.99x | 179732 | 7119546 | 77 |
| SRR28623900 | JBBFFY000000000 | SAMN40466908 | cjeju20_42 | 99.99x | 159977 | 7570538 | 76 |
| SRR28623889 | JBBFFX000000000 | SAMN40466909 | cjeju20_43 | 99.99x | 185286 | 7138718 | 233 |
| SRR28623878 | JBBFFW000000000 | SAMN40466910 | cjeju20_50 | 99.98x | 153919 | 2833076 | 83 |
| SRR28623867 | JBBFFV000000000 | SAMN40466911 | cjeju20_51 | 99.98x | 160327 | 3445572 | 43 |
| SRR28623856 | JBBFFU000000000 | SAMN40466912 | cjeju20_53 | 99.99x | 128594 | 3418962 | 91 |
| SRR28623845 | JBBFFR000000000 | SAMN40466915 | cjeju20_57 | 99.99x | 153879 | 3531346 | 66 |
| SRR28623834 | JBBFFQ000000000 | SAMN40466916 | cjeju20_59 | 99.99x | 140975 | 3024516 | 73 |
| SRR28623823 | JBBFFP000000000 | SAMN40466917 | cjeju20_60 | 99.98x | 190514 | 6592102 | 97 |
| SRR28623921 | JBBFFO000000000 | SAMN40466918 | cjeju20_61 | 99.98x | 194557 | 3283254 | 70 |
| SRR28623910 | JBBFFN000000000 | SAMN40466919 | cjeju20_62 | 99.98x | 142437 | 3363104 | 111 |
| SRR28623908 | JBBFFM000000000 | SAMN40466920 | cjeju20_8 | 99.98x | 153177 | 6740916 | 49 |
| SRR28623907 | JBBFFL000000000 | SAMN40466921 | cjeju21_1 | 99.99x | 153870 | 2188358 | 63 |
| SRR28623906 | JBBFFK000000000 | SAMN40466922 | cjeju21_13 | 99.99x | 253397 | 2411510 | 52 |
| SRR28623905 | JBBFFJ000000000 | SAMN40466923 | cjeju21_14 | 99.98x | 141357 | 2507862 | 191 |
| SRR28623904 | JBBFFI000000000 | SAMN40466924 | cjeju21_15 | 99.99x | 187004 | 2652784 | 49 |
| SRR28623903 | JBBFFH000000000 | SAMN40466925 | cjeju21_16 | 99.99x | 117866 | 2309240 | 36 |
| SRR28623902 | JBBFFG000000000 | SAMN40466926 | cjeju21_17 | 99.99x | 116389 | 2421032 | 113 |
| SRR28623901 | JBBFFF000000000 | SAMN40466927 | cjeju21_18 | 99.99x | 159865 | 2317260 | 468 |
| SRR28623899 | JBBFFE000000000 | SAMN40466928 | cjeju21_19 | 99.98x | 178288 | 2514010 | 43 |
| SRR28623898 | JBBFFD000000000 | SAMN40466929 | cjeju21_2 | 99.99x | 234121 | 2520942 | 49 |
| SRR28623897 | JBBFFC000000000 | SAMN40466930 | cjeju21_21 | 100.0x | 190041 | 2177778 | 37 |
| SRR28623896 | JBBFFC000000000 | SAMN40466931 | cjeju21_22 | 99.99x | 221390 | 1783304 | 162 |
| SRR28623895 | JBBFFA000000000 | SAMN40466932 | cjeju21_23 | 99.97xx | 336543 | 2634846 | 202 |
| SRR28623894 | JBBFEZ000000000 | SAMN40466933 | cjeju21_24 | 100.0x | 197767 | 2380728 | 34 |
| SRR28623893 | JBBFEX000000000 | SAMN40466934 | cjeju21_25 | 99.97x | 191030 | 1756604 | 54 |
| SRR28623892 | JBBFEX000000000 | SAMN40466935 | cjeju21_26 | 99.97x | 153872 | 2030236 | 52 |
| SRR28623891 | JBBFEW000000000 | SAMN40466936 | cjeju21_27 | 99.99x | 186576 | 2313994 | 212 |
| SRR28623890 | JBBFEV000000000 | SAMN40466937 | cjeju21_28 | 99.99x | 166365 | 2646550 | 75 |
| SRR28623888 | JBBFEU000000000 | SAMN40466938 | cjeju21_29 | 99.99x | 153872 | 1812620 | 55 |
| SRR28673397 | JBBFET000000000 | SAMN40466939 | cjeju21_31 | 99.98x | 179595 | 1790478 | 456 |
| SRR28623887 | JBBFES000000000 | SAMN40466940 | cjeju21_32 | 99.98x | 161836 | 2048574 | 120 |
| SRR28623886 | JBBFER000000000 | SAMN40466941 | cjeju21_33 | 99.98x | 176464 | 1659286 | 114 |
| SRR28623885 | JBBFEQ000000000 | SAMN40466942 | cjeju21_34 | 99.99x | 153903 | 2164026 | 35 |
| SRR28623884 | JBBFEP000000000 | SAMN40466943 | cjeju21_35 | 99.98x | 154432 | 2109846 | 98 |
| SRR28623883 | JBBFEO000000000 | SAMN40466944 | cjeju21_36 | 99.99x | 135602 | 1941554 | 47 |
| SRR28623882 | JBBFEN000000000 | SAMN40466945 | cjeju21_37 | 99.99x | 155765 | 1962304 | 31 |
| SRR28623881 | JBBFEM000000000 | SAMN40466946 | cjeju21_8 | 99.98x | 161702 | 1996866 | 57 |
| SRR28623880 | JBBFEL000000000 | SAMN40466947 | cjeju22_10 | 99.99x | 188013 | 2273574 | 41 |
| SRR28623879 | JBBFEK000000000 | SAMN40466948 | cjeju22_100 | 99.99x | 184212 | 2394708 | 38 |
| SRR28623877 | JBBFEJ000000000 | SAMN40466949 | cjeju22_101 | 99.99x | 153872 | 1995248 | 75 |
| SRR28623876 | JBBFEI000000000 | SAMN40466950 | cjeju22_103 | 99.99x | 213664 | 2113344 | 37 |
| SRR28623875 | JBBFEH000000000 | SAMN40466951 | cjeju22_106 | 99.98x | 195210 | 2081852 | 65 |
| SRR28623874 | JBBFEG000000000 | SAMN40466952 | cjeju22_108 | 99.98x | 185585 | 2405408 | 35 |
| SRR28623873 | JBBFEF000000000 | SAMN40466953 | cjeju22_109 | 99.98x | 157010 | 2582578 | 58 |
| SRR28623872 | JBBFEE000000000 | SAMN40466954 | cjeju22_11 | 99.99x | 129257 | 1937280 | 61 |
| SRR28623871 | JBBFED000000000 | SAMN40466955 | cjeju22_110 | 99.99x | 153881 | 2182294 | 35 |
| SRR28623870 | JBBFEC000000000 | SAMN40466956 | cjeju22_111 | 100.0x | 153903 | 2265722 | 65 |
| SRR28623869 | JBBFEB000000000 | SAMN40466957 | cjeju22_112 | 99.99x | 168232 | 2256388 | 31 |
| SRR28623868 | JBBFEA000000000 | SAMN40466958 | cjeju22_114 | 100.0x | 153872 | 1826196 | 53 |
| SRR28623866 | JBBFDZ000000000 | SAMN40466959 | cjeju22_115 | 99.97x | 220398 | 1786558 | 32 |
| SRR28623865 | JBBFDY000000000 | SAMN40466960 | cjeju22_116 | 99.99x | 334577 | 1877014 | 57 |
| SRR28623864 | JBBFDX000000000 | SAMN40466961 | cjeju22_12 | 99.99x | 113853 | 2382100 | 55 |
| SRR28623863 | JBBFDW000000000 | SAMN40466962 | cjeju22_125 | 100.0x | 186577 | 2156200 | 35 |
| SRR28623862 | JBBFDV000000000 | SAMN40466963 | cjeju22_126 | 100.0x | 106512 | 2175198 | 29 |
| SRR28623861 | JBBFDU000000000 | SAMN40466965 | cjeju22_16 | 99.99x | 153879 | 2166304 | 138 |
| SRR28623860 | JBBFDT000000000 | SAMN40466966 | cjeju22_18 | 99.98x | 298176 | 2071612 | 44 |
| SRR28623859 | JBBFDS000000000 | SAMN40466967 | cjeju22_19 | 99.98x | 188187 | 2714538 | 59 |
| SRR28623858 | JBBFDR000000000 | SAMN40466968 | cjeju22_2 | 99.99x | 177429 | 1723612 | 154 |
| SRR28623857 | JBBFDQ000000000 | SAMN40466969 | cjeju22_20 | 99.99x | 120045 | 2232882 | 38 |
| SRR28623855 | JBBFDP000000000 | SAMN40466970 | cjeju22_21 | 100.0 | 184956 | 2091988 | 49 |
| SRR28623854 | JBBFDO000000000 | SAMN40466971 | cjeju22_32 | 99.99x | 183488 | 2985470 | 102 |
| SRR28623853 | JBBFDN000000000 | SAMN40466972 | cjeju22_33 | 99.99x | 220397 | 2601094 | 81 |
| SRR28623852 | JBBFDM000000000 | SAMN40466973 | cjeju22_4 | 99.99x | 160695 | 2050346 | 70 |
| SRR28623851 | JBBFDL000000000 | SAMN40466974 | cjeju22_48 | 99.98x | 179596 | 3012910 | 35 |
| SRR28623850 | JBBFDK000000000 | SAMN40466975 | cjeju22_49 | 99.99x | 178819 | 3234746 | 1601 |
| SRR28623849 | JBBFDJ000000000 | SAMN40466976 | cjeju22_5 | 99.99x | 158338 | 2251096 | 105 |
| SRR28623848 | JBBFDI000000000 | SAMN40466977 | cjeju22_50 | 99.99x | 160665 | 2716196 | 34 |
| SRR28623847 | JBBFDH000000000 | SAMN40466978 | cjeju22_51 | 99.99x | 186576 | 2586870 | 29 |
| SRR28623846 | JBBFDG000000000 | SAMN40466979 | cjeju22_52 | 99.97x | 186595 | 3123780 | 58 |
| SRR28623844 | JBBFDF000000000 | SAMN40466980 | cjeju22_53 | 99.99x | 119380 | 2441372 | 35 |
| SRR28623843 | JBBFDE000000000 | SAMN40466981 | cjeju22_54 | 99.98x | 154154 | 3138134 | 77 |
| SRR28623842 | JBBFDD000000000 | SAMN40466982 | cjeju22_56 | 99.99x | 114939 | 2790972 | 33 |
| SRR28623841 | JBBFDC000000000 | SAMN40466983 | cjeju22_57 | 99.99x | 112646 | 2920886 | 38 |
| SRR28623840 | JBBFDB000000000 | SAMN40466984 | cjeju22_58 | 100.0x | 171801 | 2880842 | 32 |
| SRR28623839 | JBBFDA000000000 | SAMN40466985 | cjeju22_59 | 99.99x | 197766 | 2798634 | 64 |
| SRR28623838 | JBBFCZ000000000 | SAMN40466986 | cjeju22_60 | 99.99x | 159521 | 2748228 | 125 |
| SRR28623837 | JBBFCY000000000 | SAMN40466987 | cjeju22_61 | 99.99x | 334479 | 3151850 | 65 |
| SRR28623836 | JBBFCX000000000 | SAMN40466988 | cjeju22_62 | 99.99x | 178121 | 2916998 | 59 |
| SRR28623835 | JBBFCW000000000 | SAMN40466989 | cjeju22_8 | 99.99x | 148383 | 2221748 | 201 |
| SRR28623833 | JBBFCV000000000 | SAMN40466990 | cjeju22_9 | 99.99x | 174741 | 2291652 | 43 |
| SRR28623832 | JBBFCU000000000 | SAMN40466991 | cjeju22_91 | 99.98x | 214759 | 1956940 | 83 |
| SRR28623831 | JBBFCT000000000 | SAMN40466992 | cjeju22_92 | 99.99x | 186577 | 1987012 | 32 |
| SRR28673396 | JBBFCS000000000 | SAMN40466993 | cjeju22_93 | 99.99x | 152926 | 2181358 | 67 |
| SRR28623830 | JBBFCR000000000 | SAMN40466994 | cjeju22_98 | 100.0x | 160568 | 1794998 | 28 |
| SRR28623829 | JBBFCQ000000000 | SAMN40466995 | cjeju22_99 | 99.98x | 106509 | 2498484 | 27 |
| SRR28623828 | JBBFCP000000000 | SAMN40466996 | cjeju23_101 | 99.99x | 159865 | 2234146 | 38 |
| SRR28623827 | JBBFCO000000000 | SAMN40466997 | cjeju23_102 | 99.99x | 128167 | 2114884 | 51 |
| SRR28623826 | JBBFCN000000000 | SAMN40466998 | cjeju23_103 | 99.99x | 176182 | 2288428 | 109 |
| SRR28623825 | JBBFCM000000000 | SAMN40466999 | cjeju23_104 | 100.0x | 222823 | 2182680 | 31 |
| SRR28623824 | JBBFCL000000000 | SAMN40467000 | cjeju23_105 | 99.99x | 181325 | 2114520 | 70 |
| SRR28623822 | JBBFCK000000000 | SAMN40467001 | cjeju23_107 | 100.0x | 197775 | 2093676 | 40 |
| SRR28623821 | JBBFCJ000000000 | SAMN40467002 | cjeju23_108 | 99.96x | 145879 | 2101796 | 28 |
| SRR28623820 | JBBFCI000000000 | SAMN40467003 | cjeju23_60 | 99.98x | 293640 | 2003508 | 37 |
| SRR28623819 | JBBFCH000000000 | SAMN40467004 | cjeju23_63 | 99.99x | 186577 | 2211508 | 36 |
| SRR28623818 | JBBFCG000000000 | SAMN40467005 | cjeju23_65 | 99.99x | 114939 | 2036700 | 32 |
| SRR28623817 | JBBFCF000000000 | SAMN40467006 | cjeju23_67 | 99.98x | 153995 | 2140738 | 40 |
| SRR28623816 | JBBFCE000000000 | SAMN40467007 | cjeju23_68 | 99.98x | 142433 | 1973948 | 46 |
| SRR28623815 | JBBFCD000000000 | SAMN40467008 | cjeju23_83 | 99.99x | 153911 | 2154656 | 23 |
| SRR28623814 | JBBFCC000000000 | SAMN40467009 | cjeju23_84 | 99.99x | 151685 | 2237872 | 32 |
| SRR28623813 | JBBFCB000000000 | SAMN40467010 | cjeju23_89 | 100.0x | 212985 | 2529646 | 29 |
| SRR28623920 | JBBFCA000000000 | SAMN40467011 | cjeju23_91 | 99.98x | 166153 | 1885540 | 59 |
| SRR28623919 | JBBFBZ000000000 | SAMN40467012 | cjeju23_92 | 99.99x | 188160 | 2065332 | 64 |
| SRR28623918 | JBBFBY000000000 | SAMN40467013 | cjeju23_93 | 99.99x | 190186 | 3220464 | 60 |
| SRR28623917 | JBBFBX000000000 | SAMN40467014 | cjeju23_94 | 100.0x | 154048 | 2049780 | 80 |
| SRR28623916 | JBBFBW000000000 | SAMN40467015 | cjeju23_95 | 99.99x | 182518 | 2173654 | 32 |
| SRR28673396 | JBBFBV000000000 | SAMN40467016 | cjeju24_1 | 99.98x | 186721 | 2758948 | 109 |
| SRR28623915 | JBBFBU000000000 | SAMN40467017 | cjeju24_10 | 99.98x | 187571 | 2178134 | 42 |
| SRR28623914 | JBBFBT000000000 | SAMN40467018 | cjeju24_3 | 99.99x | 161015 | 2407190 | 84 |
| SRR28623913 | JBBFBS000000000 | SAMN40467019 | cjeju24_5 | 99.99x | 190060 | 2330324 | 37 |
| SRR28623912 | JBBFBR000000000 | SAMN40467020 | cjeju24_6 | 99.99x | 153184 | 2389318 | 49 |
| SRR28623911 | JBBFBQ000000000 | SAMN40467021 | cjeju24_8 | 99.99x | 186565 | 2233130 | 32 |
| SRR28623909 | JBBFBP000000000 | SAMN40467022 | cjeju24_9 | 99.99x | 186577 | 2432328 | 68 |

**Table S2.**
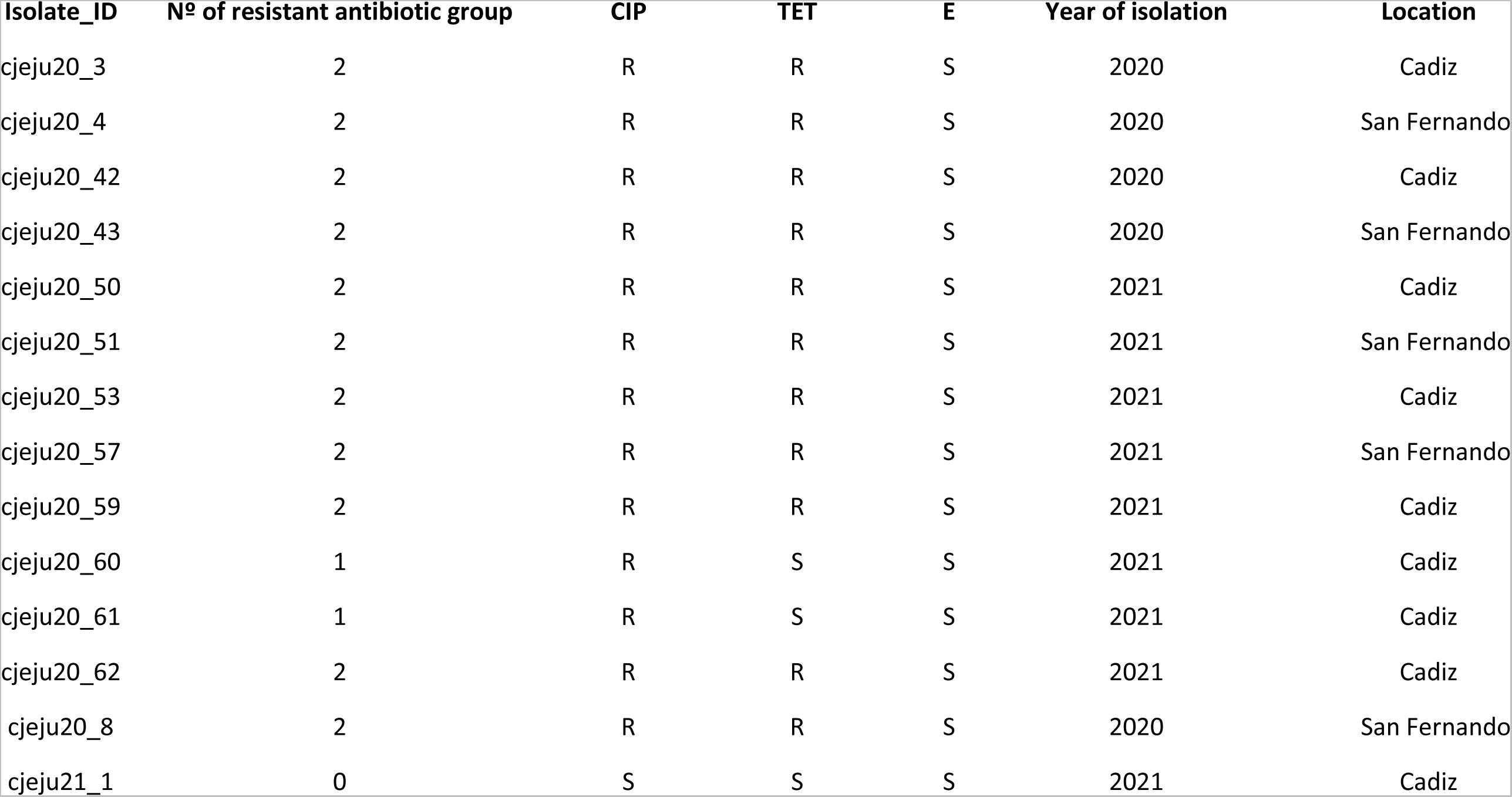
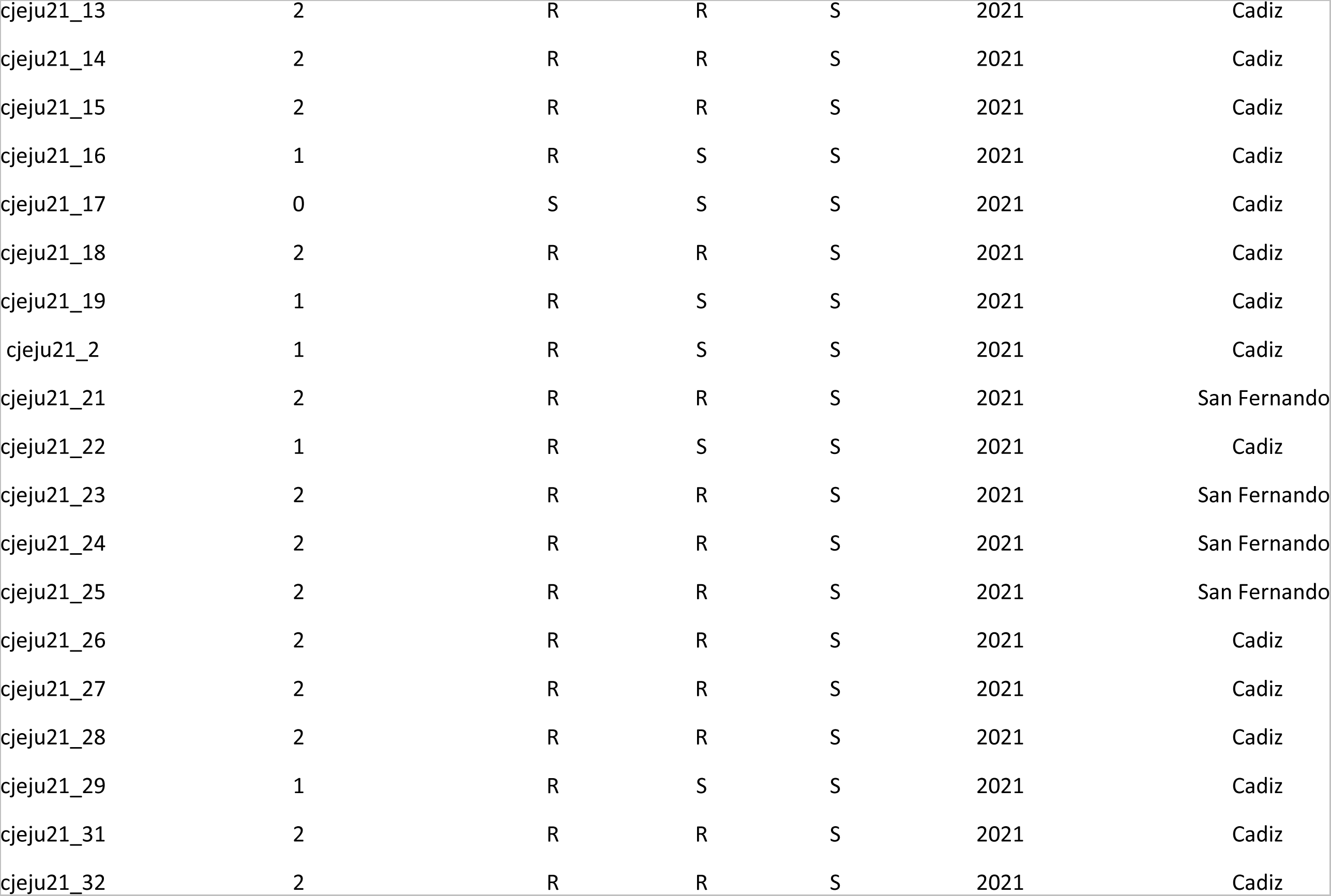
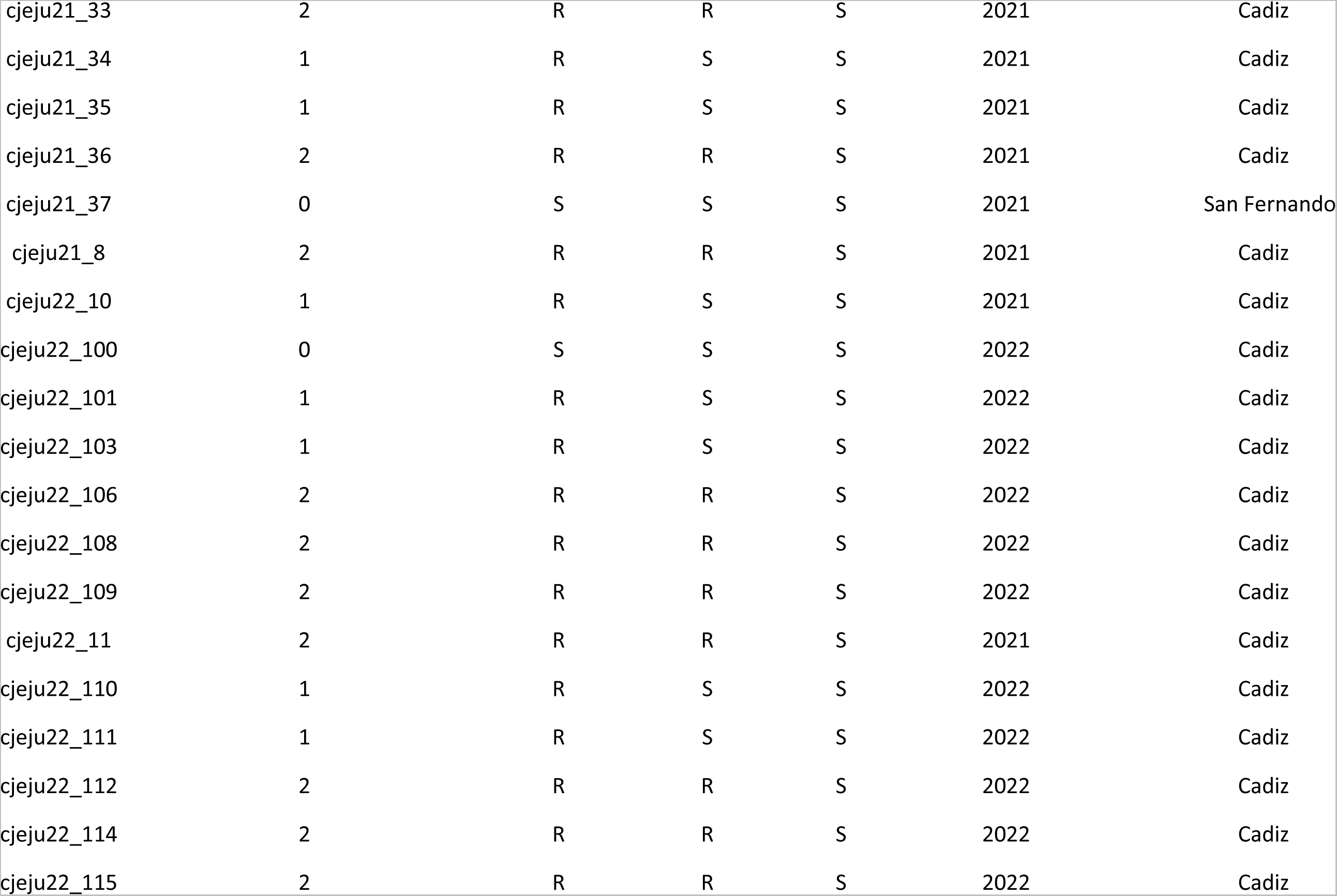
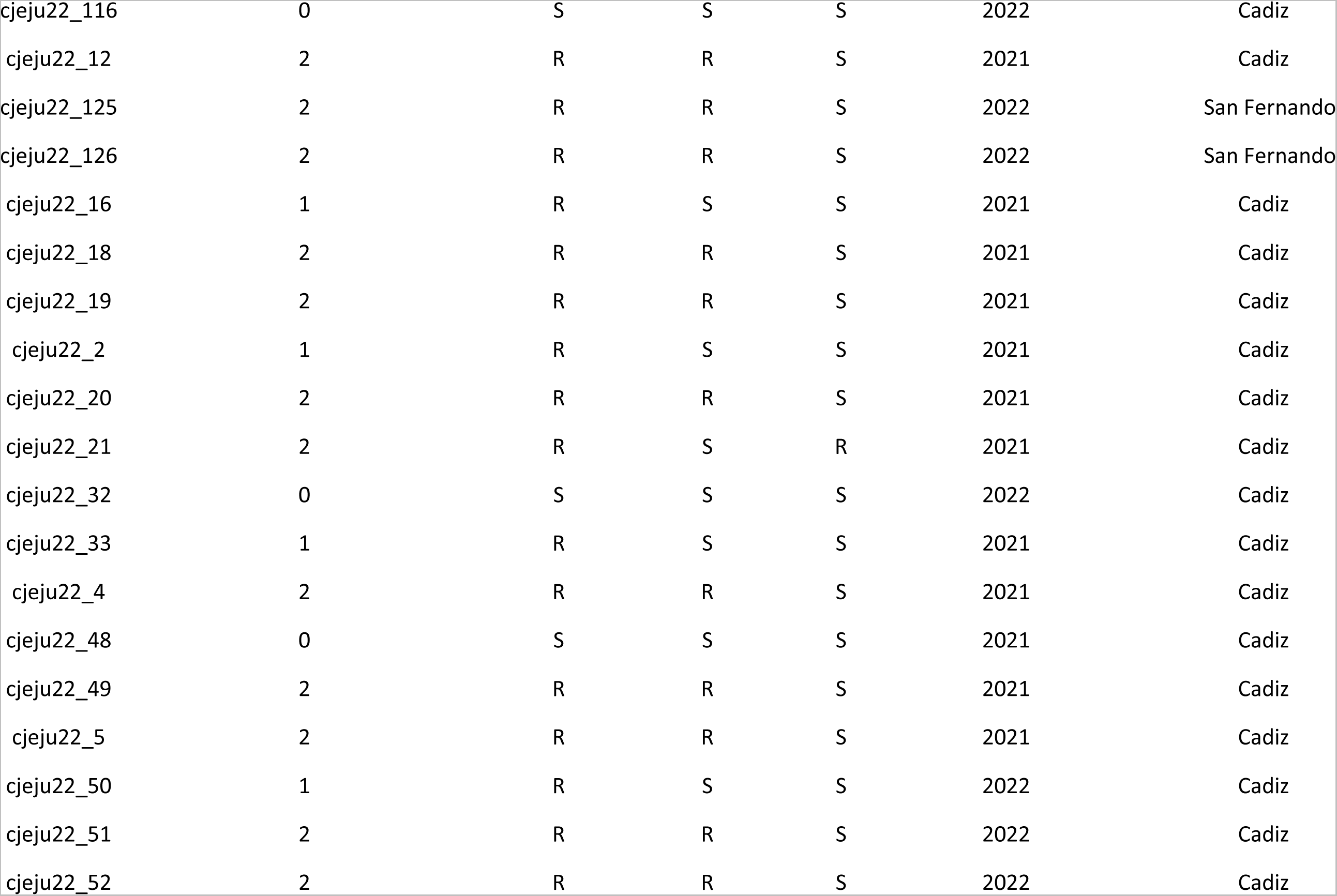
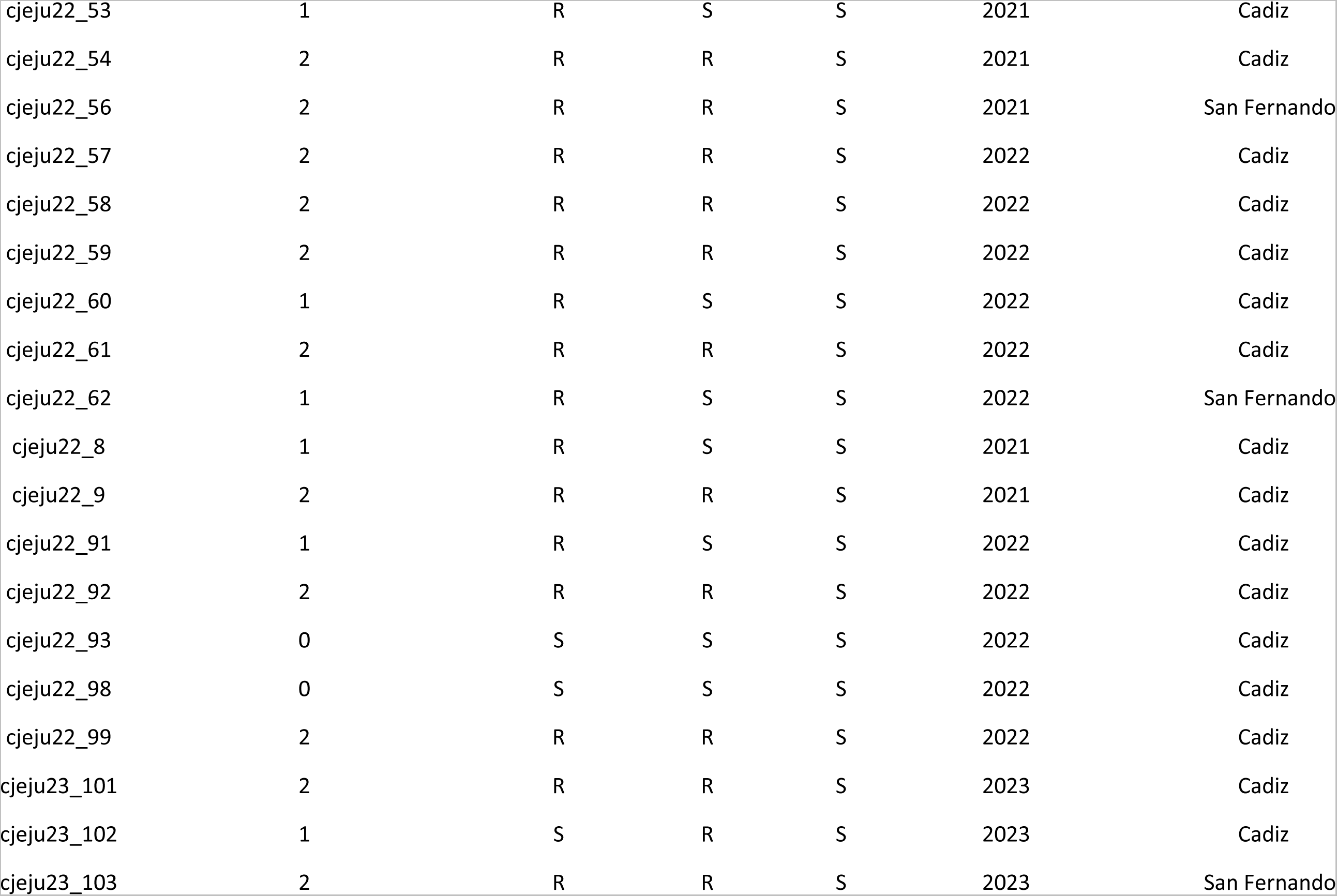
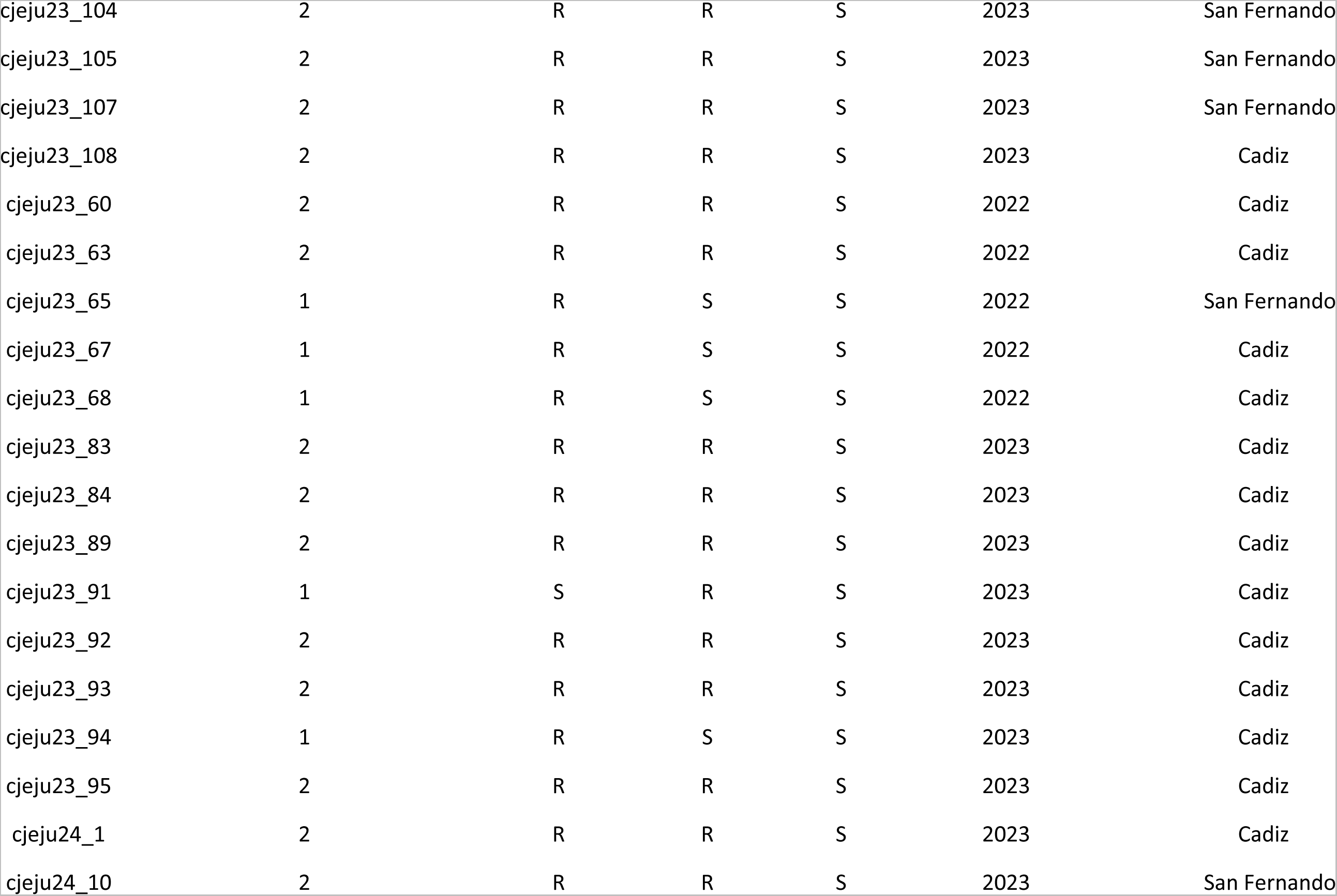
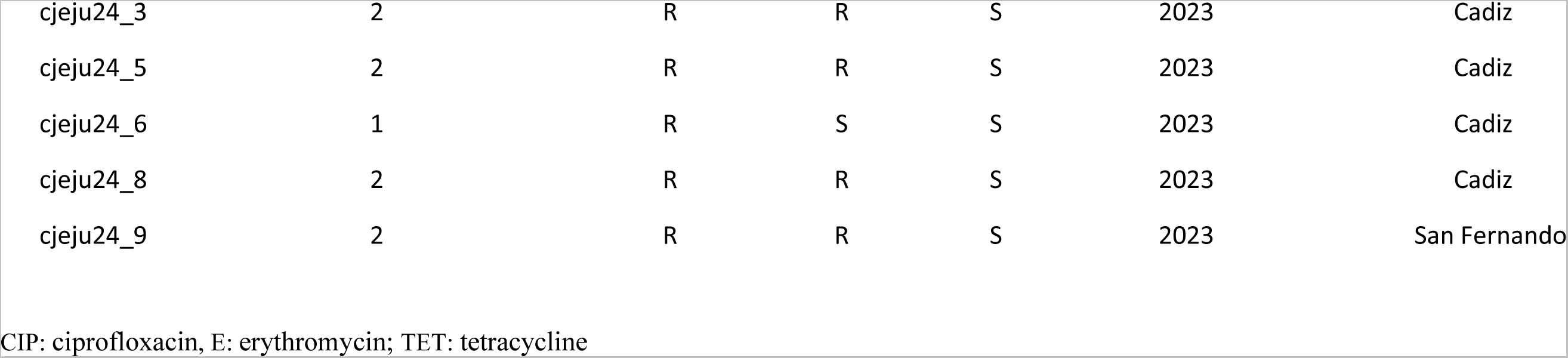
Antimicrobial susceptibility, number of resistant antibiotic group, years of isolation and location of 114 *Campylobacter jejuni* isolates.

| Isolate_ID | Nº of resistant antibiotic group | CIP | TET | E | Year of isolation | Location |
| --- | --- | --- | --- | --- | --- | --- |
| cjeju20_3 | 2 | R | R | S | 2020 | Cadiz |
| cjeju20_4 | 2 | R | R | S | 2020 | San Fernando |
| cjeju20_42 | 2 | R | R | S | 2020 | Cadiz |
| cjeju20_43 | 2 | R | R | S | 2020 | San Fernando |
| cjeju20_50 | 2 | R | R | S | 2021 | Cadiz |
| cjeju20_51 | 2 | R | R | S | 2021 | San Fernando |
| cjeju20_53 | 2 | R | R | S | 2021 | Cadiz |
| cjeju20_57 | 2 | R | R | S | 2021 | San Fernando |
| cjeju20_59 | 2 | R | R | S | 2021 | Cadiz |
| cjeju20_60 | 1 | R | S | S | 2021 | Cadiz |
| cjeju20_61 | 1 | R | S | S | 2021 | Cadiz |
| cjeju20_62 | 2 | R | R | S | 2021 | Cadiz |
| cjeju20_8 | 2 | R | R | S | 2020 | San Fernando |
| cjeju21_1 | 0 | S | S | S | 2021 | Cadiz |
| cjeju21_13 | 2 | R | R | S | 2021 | Cadiz |
| cjeju21_14 | 2 | R | R | S | 2021 | Cadiz |
| cjeju21_15 | 2 | R | R | S | 2021 | Cadiz |
| cjeju21_16 | 1 | R | S | S | 2021 | Cadiz |
| cjeju21_17 | 0 | S | S | S | 2021 | Cadiz |
| cjeju21_18 | 2 | R | R | S | 2021 | Cadiz |
| cjeju21_19 | 1 | R | S | S | 2021 | Cadiz |
| cjeju21_2 | 1 | R | S | S | 2021 | Cadiz |
| cjeju21_21 | 2 | R | R | S | 2021 | San Fernando |
| cjeju21_22 | 1 | R | S | S | 2021 | Cadiz |
| cjeju21_23 | 2 | R | R | S | 2021 | San Fernando |
| cjeju21_24 | 2 | R | R | S | 2021 | San Fernando |
| cjeju21_25 | 2 | R | R | S | 2021 | San Fernando |
| cjeju21_26 | 2 | R | R | S | 2021 | Cadiz |
| cjeju21_27 | 2 | R | R | S | 2021 | Cadiz |
| cjeju21_28 | 2 | R | R | S | 2021 | Cadiz |
| cjeju21_29 | 1 | R | S | S | 2021 | Cadiz |
| cjeju21_31 | 2 | R | R | S | 2021 | Cadiz |
| cjeju21_32 | 2 | R | R | S | 2021 | Cadiz |
| cjeju21_33 | 2 | R | R | S | 2021 | Cadiz |
| cjeju21_34 | 1 | R | S | S | 2021 | Cadiz |
| cjeju21_35 | 1 | R | S | S | 2021 | Cadiz |
| cjeju21_36 | 2 | R | R | S | 2021 | Cadiz |
| cjeju21_37 | 0 | S | S | S | 2021 | San Fernando |
| cjeju21_8 | 2 | R | R | S | 2021 | Cadiz |
| cjeju22_10 | 1 | R | S | S | 2021 | Cadiz |
| cjeju22_100 | 0 | S | S | S | 2022 | Cadiz |
| cjeju22_101 | 1 | R | S | S | 2022 | Cadiz |
| cjeju22_103 | 1 | R | S | S | 2022 | Cadiz |
| cjeju22_106 | 2 | R | R | S | 2022 | Cadiz |
| cjeju22_108 | 2 | R | R | S | 2022 | Cadiz |
| cjeju22_109 | 2 | R | R | S | 2022 | Cadiz |
| cjeju22_11 | 2 | R | R | S | 2021 | Cadiz |
| cjeju22_110 | 1 | R | S | S | 2022 | Cadiz |
| cjeju22_111 | 1 | R | S | S | 2022 | Cadiz |
| cjeju22_112 | 2 | R | R | S | 2022 | Cadiz |
| cjeju22_114 | 2 | R | R | S | 2022 | Cadiz |
| cjeju22_115 | 2 | R | R | S | 2022 | Cadiz |
| cjeju22_116 | 0 | S | S | S | 2022 | Cadiz |
| cjeju22_12 | 2 | R | R | S | 2021 | Cadiz |
| cjeju22_125 | 2 | R | R | S | 2022 | San Fernando |
| cjeju22_126 | 2 | R | R | S | 2022 | San Fernando |
| cjeju22_16 | 1 | R | S | S | 2021 | Cadiz |
| cjeju22_18 | 2 | R | R | S | 2021 | Cadiz |
| cjeju22_19 | 2 | R | R | S | 2021 | Cadiz |
| cjeju22_2 | 1 | R | S | S | 2021 | Cadiz |
| cjeju22_20 | 2 | R | R | S | 2021 | Cadiz |
| cjeju22_21 | 2 | R | S | R | 2021 | Cadiz |
| cjeju22_32 | 0 | S | S | S | 2022 | Cadiz |
| cjeju22_33 | 1 | R | S | S | 2021 | Cadiz |
| cjeju22_4 | 2 | R | R | S | 2021 | Cadiz |
| cjeju22_48 | 0 | S | S | S | 2021 | Cadiz |
| cjeju22_49 | 2 | R | R | S | 2021 | Cadiz |
| cjeju22_5 | 2 | R | R | S | 2021 | Cadiz |
| cjeju22_50 | 1 | R | S | S | 2022 | Cadiz |
| cjeju22_51 | 2 | R | R | S | 2022 | Cadiz |
| cjeju22_52 | 2 | R | R | S | 2022 | Cadiz |
| cjeju22_53 | 1 | R | S | S | 2021 | Cadiz |
| cjeju22_54 | 2 | R | R | S | 2021 | Cadiz |
| cjeju22_56 | 2 | R | R | S | 2021 | San Fernando |
| cjeju22_57 | 2 | R | R | S | 2022 | Cadiz |
| cjeju22_58 | 2 | R | R | S | 2022 | Cadiz |
| cjeju22_59 | 2 | R | R | S | 2022 | Cadiz |
| cjeju22_60 | 1 | R | S | S | 2022 | Cadiz |
| cjeju22_61 | 2 | R | R | S | 2022 | Cadiz |
| cjeju22_62 | 1 | R | S | S | 2022 | San Fernando |
| cjeju22_8 | 1 | R | S | S | 2021 | Cadiz |
| cjeju22_9 | 2 | R | R | S | 2021 | Cadiz |
| cjeju22_91 | 1 | R | S | S | 2022 | Cadiz |
| cjeju22_92 | 2 | R | R | S | 2022 | Cadiz |
| cjeju22_93 | 0 | S | S | S | 2022 | Cadiz |
| cjeju22_98 | 0 | S | S | S | 2022 | Cadiz |
| cjeju22_99 | 2 | R | R | S | 2022 | Cadiz |
| cjeju23_101 | 2 | R | R | S | 2023 | Cadiz |
| cjeju23_102 | 1 | S | R | S | 2023 | Cadiz |
| cjeju23_103 | 2 | R | R | S | 2023 | San Fernando |
| cjeju23_104 | 2 | R | R | S | 2023 | San Fernando |
| cjeju23_105 | 2 | R | R | S | 2023 | San Fernando |
| cjeju23_107 | 2 | R | R | S | 2023 | San Fernando |
| cjeju23_108 | 2 | R | R | S | 2023 | Cadiz |
| cjeju23_60 | 2 | R | R | S | 2022 | Cadiz |
| cjeju23_63 | 2 | R | R | S | 2022 | Cadiz |
| cjeju23_65 | 1 | R | S | S | 2022 | San Fernando |
| cjeju23_67 | 1 | R | S | S | 2022 | Cadiz |
| cjeju23_68 | 1 | R | S | S | 2022 | Cadiz |
| cjeju23_83 | 2 | R | R | S | 2023 | Cadiz |
| cjeju23_84 | 2 | R | R | S | 2023 | Cadiz |
| cjeju23_89 | 2 | R | R | S | 2023 | Cadiz |
| cjeju23_91 | 1 | S | R | S | 2023 | Cadiz |
| cjeju23_92 | 2 | R | R | S | 2023 | Cadiz |
| cjeju23_93 | 2 | R | R | S | 2023 | Cadiz |
| cjeju23_94 | 1 | R | S | S | 2023 | Cadiz |
| cjeju23_95 | 2 | R | R | S | 2023 | Cadiz |
| cjeju24_1 | 2 | R | R | S | 2023 | Cadiz |
| cjeju24_10 | 2 | R | R | S | 2023 | San Fernando |
| cjeju24_3 | 2 | R | R | S | 2023 | Cadiz |
| cjeju24_5 | 2 | R | R | S | 2023 | Cadiz |
| cjeju24_6 | 1 | R | S | S | 2023 | Cadiz |
| cjeju24_8 | 2 | R | R | S | 2023 | Cadiz |
| cjeju24_9 | 2 | R | R | S | 2023 | San Fernando |

CIP: ciprofloxacin, E: erythromycin; TET: tetracycline

**Table S3.** Distribution of Clonal complex (CC), Sequnce Type (ST) and Core genome sequence type (cgMLST) on 114 *Campylobacter jejuni* isolates.

| Isolate ID | Clonal complex (CC) | Sequence type (ST) | Core genome (cgMLST) |
| --- | --- | --- | --- |
| cjeju20_3 | ST-42 complex | 6532 | 19242 |
| cjeju20_4 | ST-42 complex | 6532 | 35104 |
| cjeju20_42 | ST-42 complex | 6532 | 14959 |
| cjeju20_43 | ST-21 complex | 50 | 88265 |
| cjeju20_50 | ST-607 complex | 607 | 45535 |
| cjeju20_51 | ST-42 complex | 6532 | 14339, 19242 |
| cjeju20_53 | ST-354 complex | 354 | 48132 |
| cjeju20_57 | ST-21 complex | 19 | 48269 |
| cjeju20_59 | ST-443 complex | 6522 | 3286 |
| cjeju20_60 | ST-353 complex | 5 | 75401 |
| cjeju20_61 | ST-443 complex | 7517 | 20116 |
| cjeju20_62 | ST-42 complex | 6532 | 19242 |
| cjeju20_8 | ST-21 complex | 50 | 19078, 81895 |
| cjeju21_1 | ST-22 complex | 22 | 81330 |
| cjeju21_13 | ST-21 complex | 50 | 25897, 88265 |
| cjeju21_14 | ST-206 complex | 572 | 79973 |
| cjeju21_15 | ST-42 complex | 6532 | 19242 |
| cjeju21_16 | ST-443 complex | 7517 | 21997 |
| cjeju21_17 | ST-22 complex | 22 | 79379 |
| cjeju21_18 | ST-48 complex | 48 | 75718 |
| cjeju21_19 | ST-52 complex | 52 | 48512 |
| cjeju21_2 | ST-21 complex | 19 | 3520 |
| cjeju21_21 | ST-257 complex | 990 | 69637 |
| cjeju21_22 | ST-443 complex | 7517 | 21997 |
| cjeju21_23 | ST-353 complex | 400 | 35312, 45012 |
| cjeju21_24 | ST-21 complex | 50 | 3006 |
| cjeju21_25 | ST-206 complex | 122 | 48438 |
| cjeju21_26 | ST-21 complex | 50 | 18998, 81843 |
| cjeju21_27 | ST-574 complex | 3172 | 74312 |
| cjeju21_28 | ST-42 complex | 6532 | 19242 |
| cjeju21_29 | ST-354 complex | 9203 | 16975, 45360, 80470 |
| cjeju21_31 | ST-607 complex | 607 | 84646 |
| cjeju21_32 | ST-353 complex | 9887 | 20782 |
| cjeju21_33 | - | 531 | 101138 |
| cjeju21_34 | ST-353 complex | 2364 | 15041 |
| cjeju21_35 | ST-21 complex | 50 | 18726,81657 |
| cjeju21_36 | ST-48 complex | 48 | 75718 |
| cjeju21_37 | - | 1080 | 76297 |
| cjeju21_8 | ST-21 complex | 50 | 79730 |
| cjeju22_10 | ST-42 complex | 6532 | 35104 |
| cjeju22_100 | ST-45 complex | 583 | 71873 |
| cjeju22_101 | ST-353 complex | 5 | 75401 |
| cjeju22_103 | ST-206 complex | 122 | 3580 |
| cjeju22_106 | ST-257 complex | 824 | 80478 |
| cjeju22_108 | ST-206 complex | 572 | 85207 |
| cjeju22_109 | ST-21 complex | 982 | 18802 |
| cjeju22_11 | ST-21 complex | 50 | 18998, 81843 |
| cjeju22_110 | ST-21 complex | 19 | 70614 |
| cjeju22_111 | ST-443 complex | 7517 | 20116, 21997 |
| cjeju22_112 | ST-206 complex | 122 | 35432 |
| cjeju22_114 | ST-354 complex | 354 | 3662 |
| cjeju22_115 | ST-206 complex | 572 | 78144 |
| cjeju22_116 | ST-257 complex | 824 | 8646 |
| cjeju22_12 | ST-574 complex | 3172 | 74312 |
| cjeju22_125 | ST-42 complex | 6532 | 14339 |
| cjeju22_126 | ST-206 complex | 572 | 78144 |
| cjeju22_16 | ST-354 complex | 9203 | 80470 |
| cjeju22_18 | ST-354 complex | 354 | 3662 |
| cjeju22_19 | ST-21 complex | 21 | 21464 |
| cjeju22_2 | ST-21 complex | 19 | 3520 |
| cjeju22_20 | Unassigend | 2133 | 72177 |
| cjeju22_21 | ST-42 complex | 6532 | 35104 |
| cjeju22_32 | ST-61 complex | 61 | 102335 |
| cjeju22_33 | ST-353 complex | 353 | 68510 |
| cjeju22_4 | ST-206 complex | 572 | 79973 |
| cjeju22_48 | ST-45 complex | 583 | 68071 |
| cjeju22_49 | ST-353 complex | 7355 | 34814, 35324 |
| cjeju22_5 | ST-21 complex | 21 | 580 |
| cjeju22_50 | ST-52 complex | 52 | 48512 |
| cjeju22_51 | ST-206 complex | 572 | 85207 |
| cjeju22_52 | ST-21 complex | 50 | 101149 |
| cjeju22_53 | ST-42 complex | 6532 | 35488 |
| cjeju22_54 | ST-257 complex | 824 | 80478 |
| cjeju22_56 | ST-353 complex | 356 | 13832 |
| cjeju22_57 | Unassigend | 2133 | 72177 |
| cjeju22_58 | ST-206 complex | 572 | 78144 |
| cjeju22_59 | ST-354 complex | 354 | 3662 |
| cjeju22_60 | ST-21 complex | 148 | 19245 |
| cjeju22_61 | ST-206 complex | 572 | 101146 |
| cjeju22_62 | ST-22 complex | 22 | 70553 |
| cjeju22_8 | ST-21 complex | 19 | 3520 |
| cjeju22_9 | ST-42 complex | 6532 | 35104 |
| cjeju22_91 | ST-353 complex | 6461 | 4628 |
| cjeju22_92 | ST-353 complex | 356 | 13832 |
| cjeju22_93 | ST-179 complex | 4447 | 99837, 99838 |
| cjeju22_98 | Unassignd | 2861 | 78144, 18045, 18046, 18047, 50989, 50990, 50991, 81273, 81274, 81275 |
| cjeju22_99 | ST-206 complex | 572 | 78144 |
| cjeju23_101 | ST-206 complex | 572 | 85207 |
| cjeju23_102 | ST-21 complex | 19 | 19368 |
| cjeju23_103 | ST-353 complex | 6461 | 4628 |
| cjeju23_104 | ST-48 complex | 48 | 77195 |
| cjeju23_105 | ST-607 complex | 607 | 4193 |
| cjeju23_107 | ST-206 complex | 122 | 35260 |
| cjeju23_108 | ST-206 complex | 572 | 85207 |
| cjeju23_60 | ST-42 complex | 517 | 35488 |
| cjeju23_63 | ST-48 complex | 48 | 75718 |
| cjeju23_65 | ST-52 complex | 52 | 48465 |
| cjeju23_67 | ST-443 complex | 7517 | 21997 |
| cjeju23_68 | ST-443 complex | 51 | 19911 |
| cjeju23_83 | ST-257 complex | 2254 | 18653 |
| cjeju23_84 | ST-257 complex | 2254 | 18653 |
| cjeju23_89 | ST-21 complex | 50 | 3006 |
| cjeju23_91 | ST-21 complex | 19 | 19368 |
| cjeju23_92 | ST-353 complex | 1232 | 20645 |
| cjeju23_93 | ST-353 complex | 8334 | 35609 |
| cjeju23_94 | ST-21 complex | 861 | 19143 |
| cjeju23_95 | ST-354 complex | 354 | 47011 |
| cjeju24_1 | ST-21 complex | 148 | 19245 |
| cjeju24_10 | ST-22 complex | 22 | 100524 |
| cjeju24_3 | ST-353 complex | 9390 | 45948 |
| cjeju24_5 | ST-206 complex | 572 | 101146 |
| cjeju24_6 | ST-45 complex | 418 | 73957 |
| cjeju24_8 | ST-206 complex | 572 | 101146 |
| cjeju24_9 | ST-354 complex | 354 | 5005 |

**Table S4.**
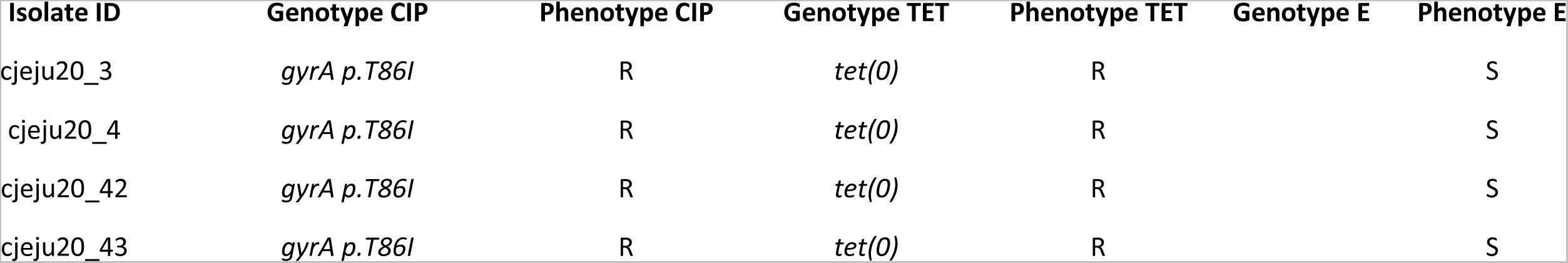
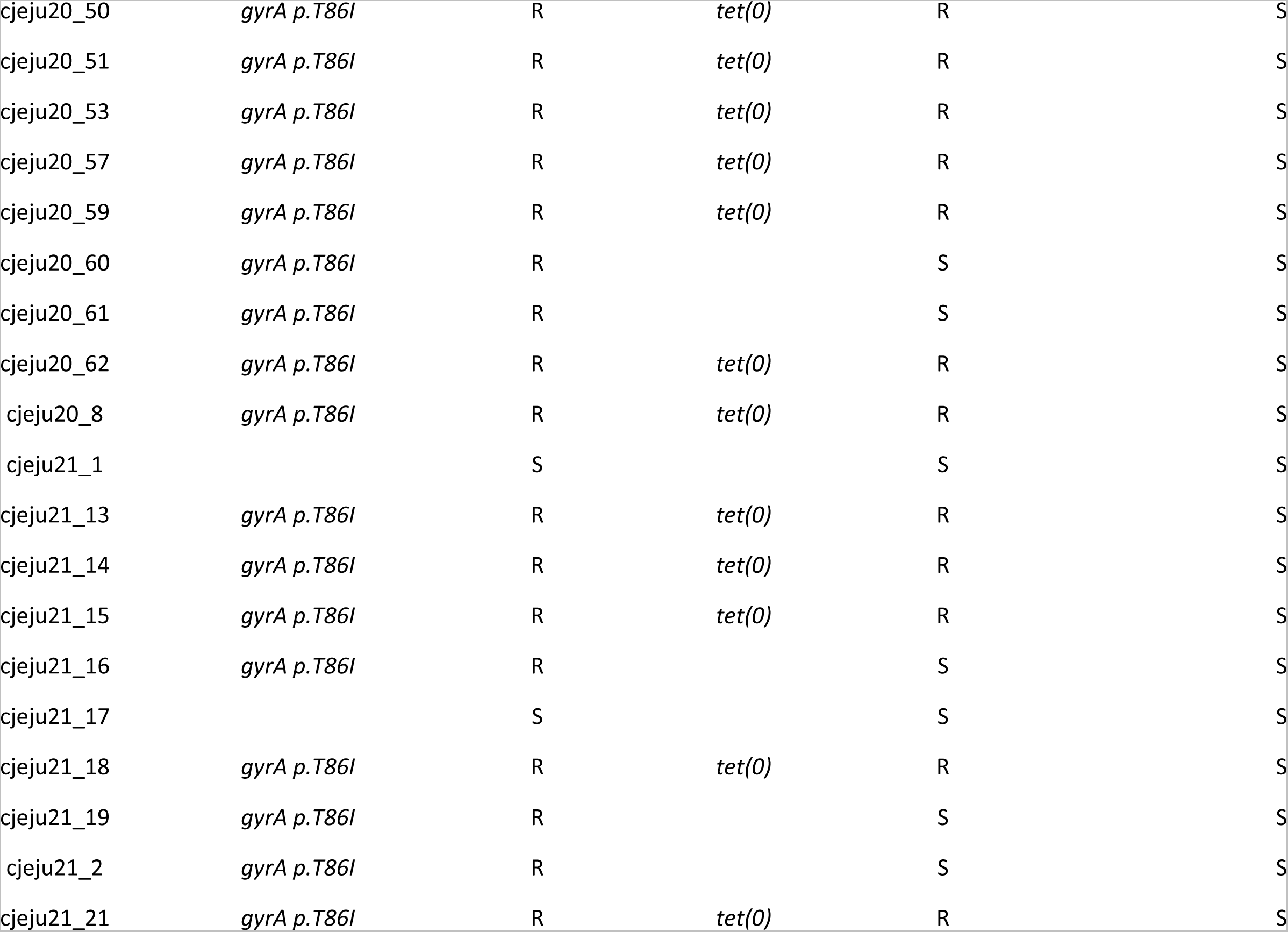
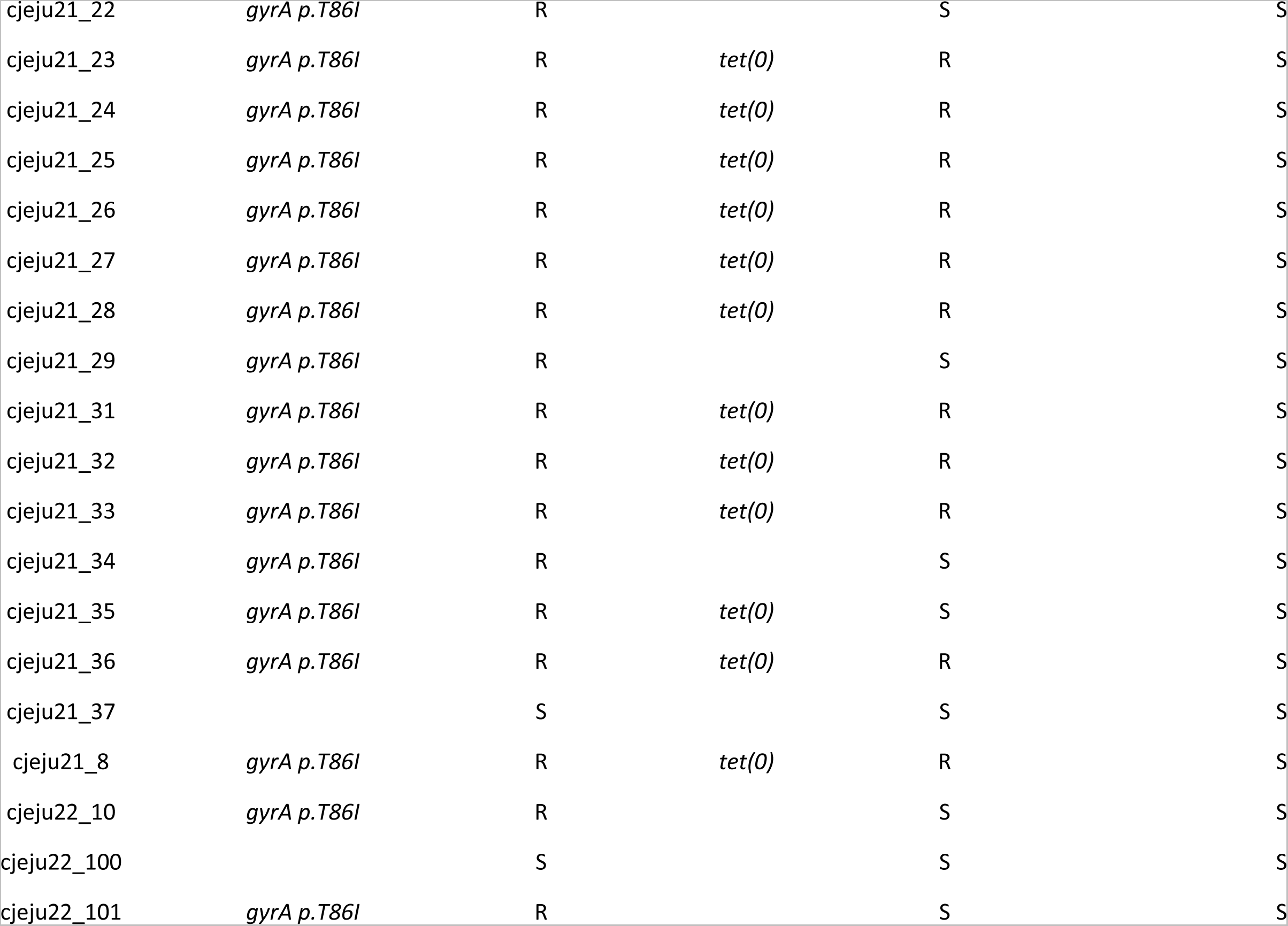
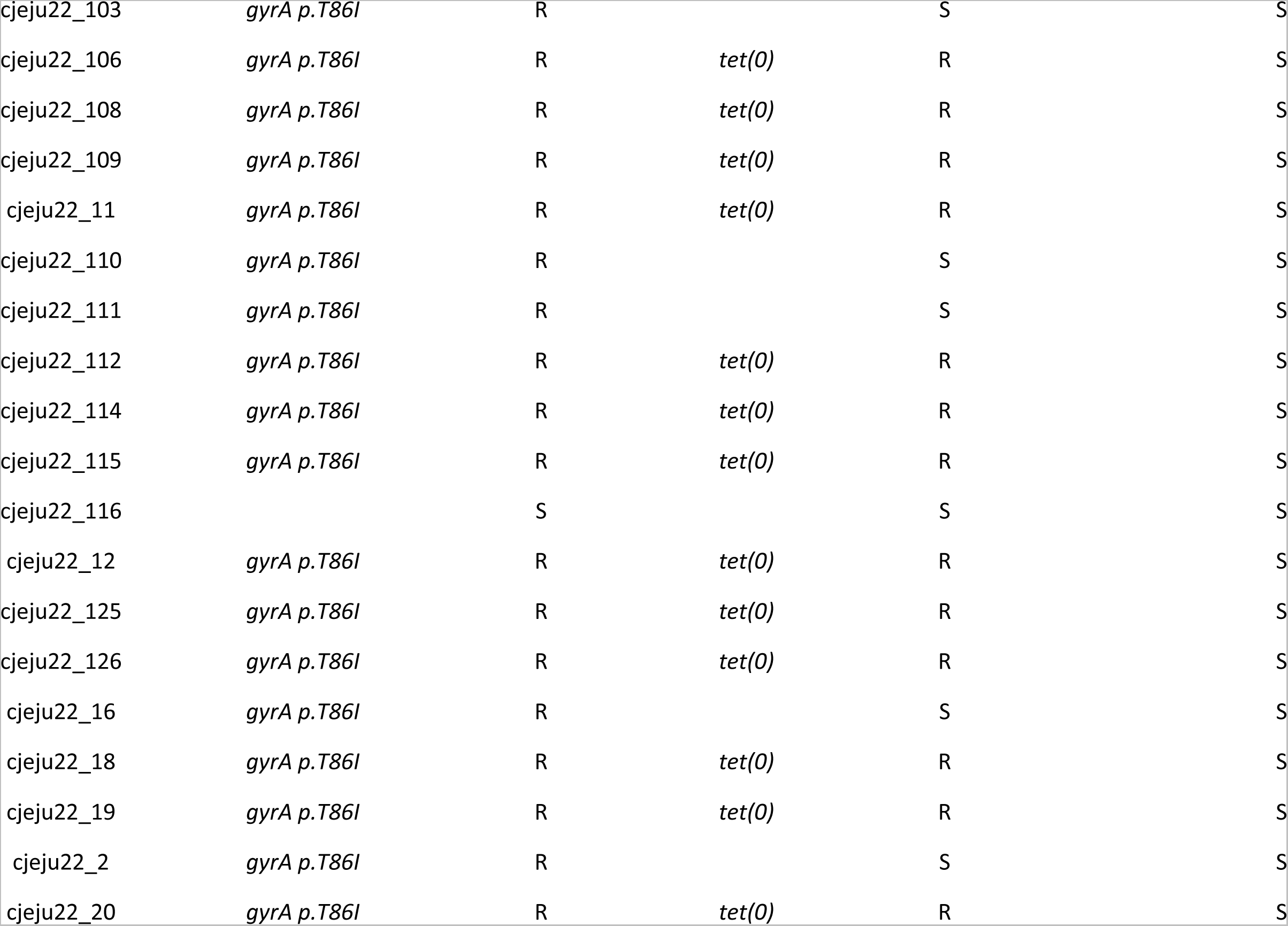
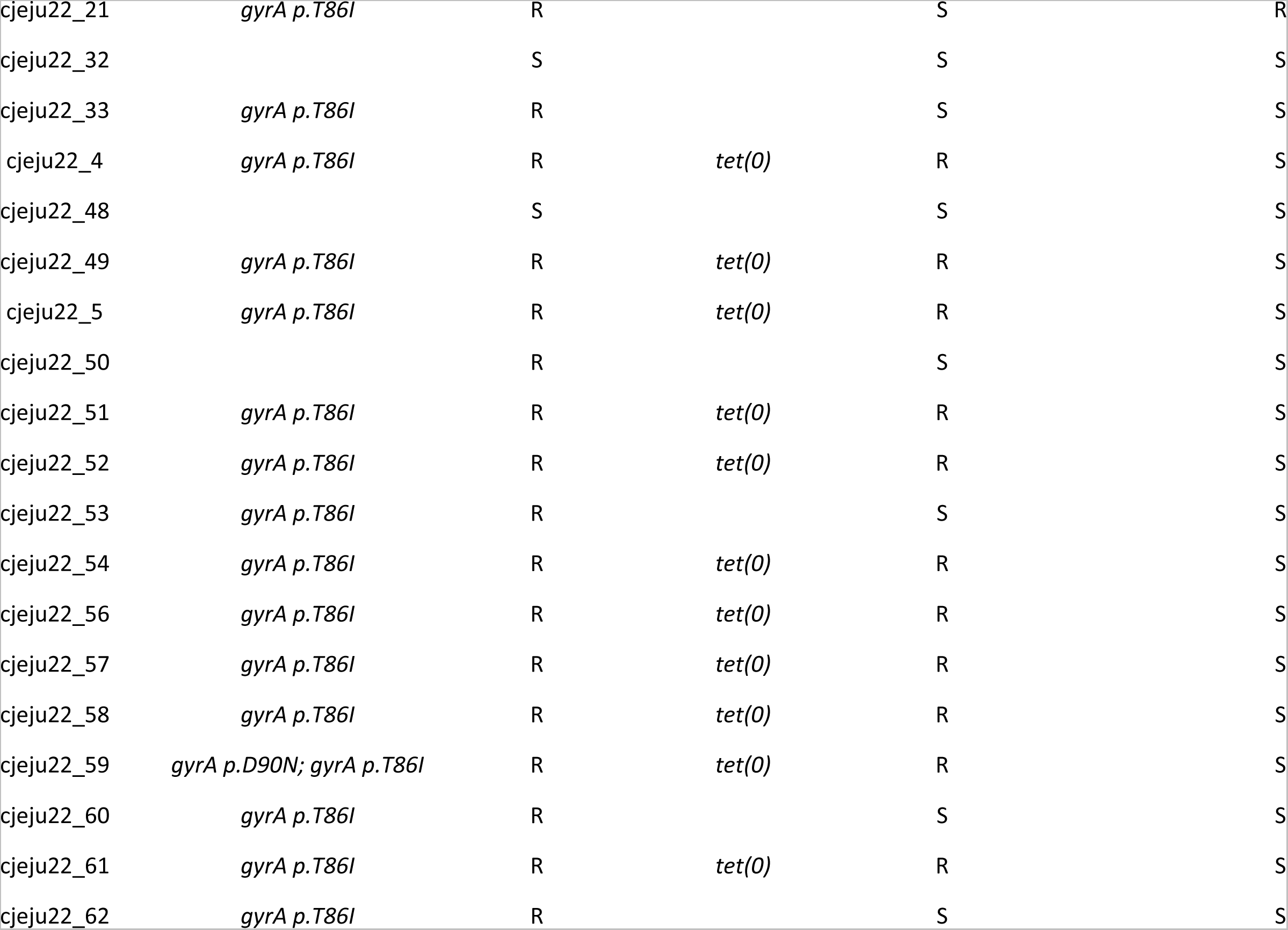
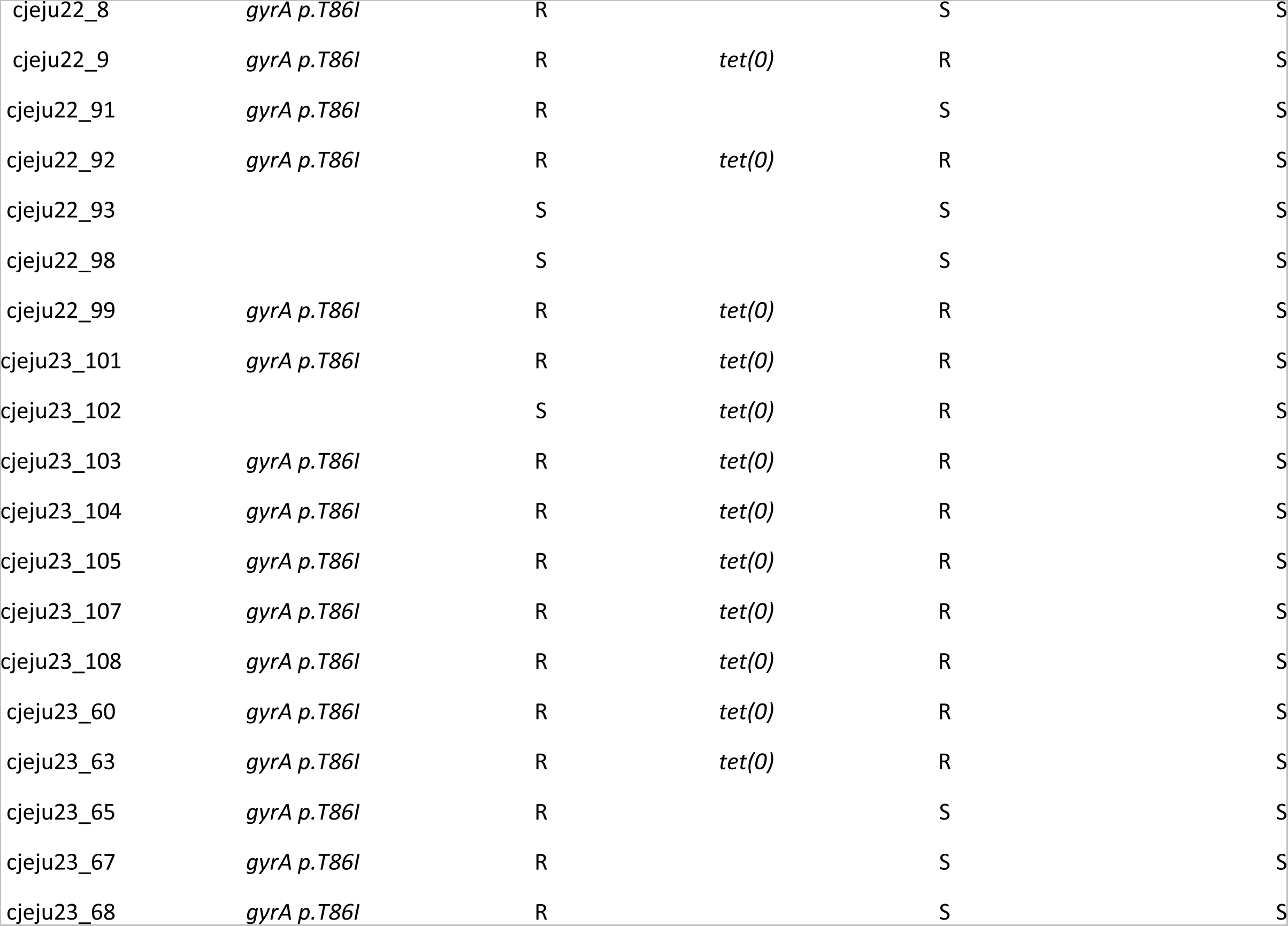
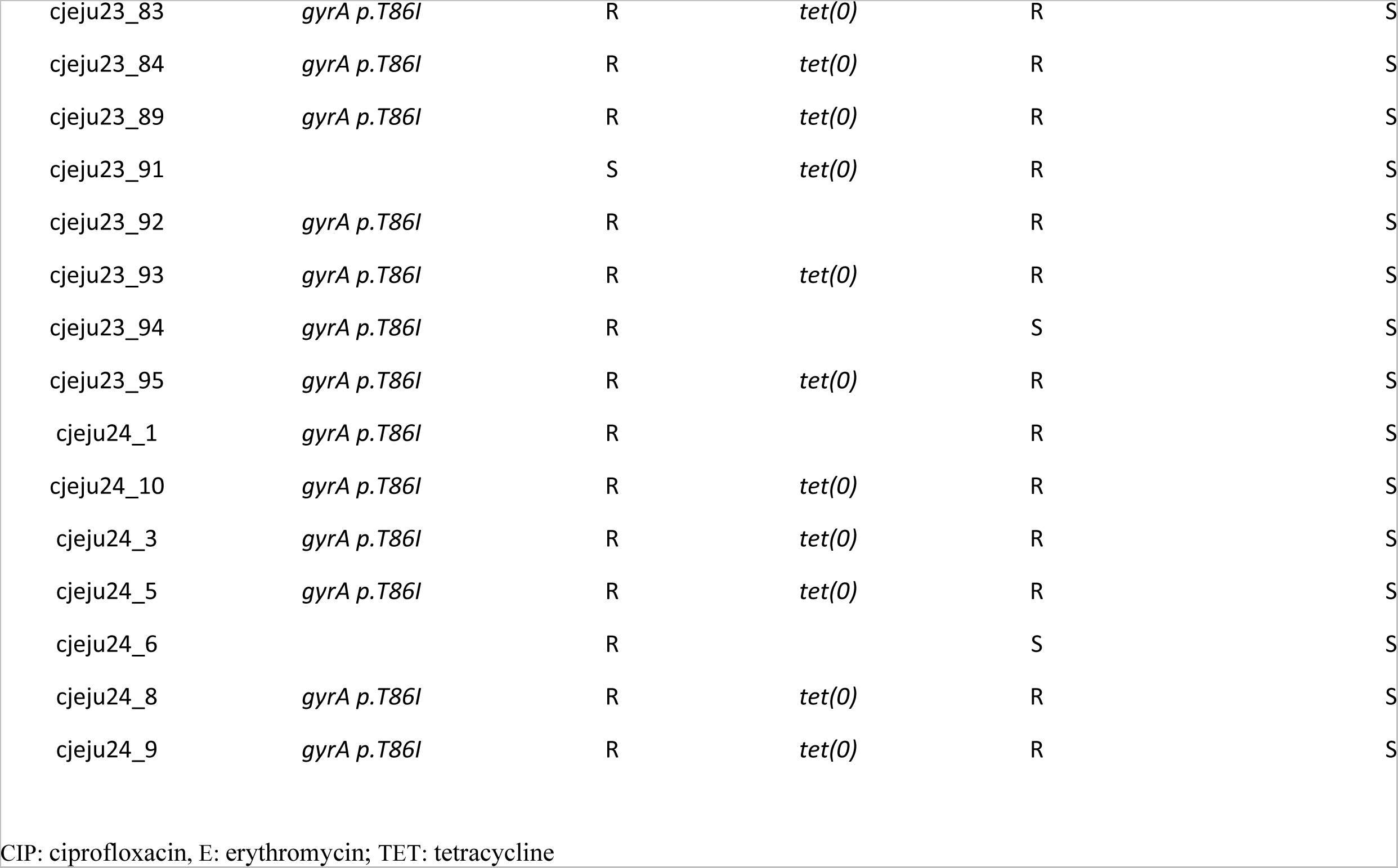
Phenotypic and genotype profiles of 114 *Campylobacter jejuni* isolates.

